# A network approach with motion sequencing reveals hidden patterns of repetitive behavior in a pre-clinical model of epilepsy

**DOI:** 10.64898/2025.12.04.692371

**Authors:** Jennifer L. Koehler, Quinn R. Harvey, Olivia R. Hoffman, Jose Ezekiel Clemente Espina, Barry A. Schoenike, Cameron P.E. Jones, Grant L. Weiss, Jamie L. Maguire, Avtar S. Roopra

## Abstract

Epilepsy is the 4th most prevalent neurological condition with 50 million cases worldwide. Patients with epilepsy bare a disproportionate burden of cognitive decline and psychiatric disorders which remain poorly understood and go unaddressed by current anti-epileptic treatments. Furthermore, pre-clinical work on behavioral comorbidities can be hampered by current testing frameworks which rely on well-defined, discreet tests with limited repeatability. Recent work has demonstrated a role for machine learning modalities such as Motion Sequencing (MoSeq) in assessing behavioral differences between naive and epileptic. In this study we combined MoSeq with a novel analysis pipeline to uncover repetitive behaviors in chronically epileptic mice. These repetitive behaviors emerge alongside epilepsy specific racing behaviors which persist in epileptic mice as disease progresses. We show that epileptic mice have more fragile and dispersed behavioral networks. Finally, we test this pipeline using the FDA approved anti-seizure medication carbamazepine, showing a rescue of racing syllable and a partial rescue of behavioral network dispersion. Together, these results lay a groundwork for extracting clinically relevant phenotypes from MoSeq data throughout disease progression.

## Introduction

Epilepsy is the 4^th^ most common neurological condition affecting over 65 million people worldwide(1). Beyond seizures, more than 50% of people with epilepsy experience at least one psychiatric comorbidity(2,3) including depression, anxiety, obsessive compulsive disorder (OCD), and attention deficit hyperactivity disorder (ADHD)(4–9). These comorbidities result in reduced quality of life and often go unaddressed by current anti-seizure medications(6,10–12). There is a lack of consensus regarding the relationship between behavioral comorbidities and seizures. One school of thought proposes that seizures drive the emergence of comorbidities whereas another hypothesizes that an underlying mechanism drives both(2,13,14).

Having psychiatric comorbidities is associated with both more severe epilepsy and drug refractory disease(15,16). The mechanism driving this association is largely unknown(17). However, there are several factors associated with both psychiatric disorders and epilepsy including but not limited to: hypothalamic-pituitary-adrenal (HPA) axis dysfunction, dysregulated neurotransmitters, and inflammatory mechanisms(2,18).

Current tools available to analyze behavioral comorbidities rely on discrete tests, such as the marble burying test for compulsive burying and Go-no Go test for impulsive behavior, which have well defined but limited outcome measures. Furthermore, certain behavioral tests risk carry-over effects when repeated too often in close succession(20). In contrast, Motion Sequencing (MoSeq), a machine learning platform combined with 3D video analysis, finds behavioral differences between epileptic and non-epileptic mice at a frame-by-frame rate(30–34) in an unbiased manner. MoSeq divides behavior into sub-second modules of behavior, known as “syllables,” comprised of activities such as a “scrunch” or “rear-up”. Others have used rates of syllable usage to distinguish epileptic and naïve mice as well as identify seizure behavior during interictal periods(25). However, while each syllable on its own has little clinical relevance, we reason that when a sequence of syllables is combined in chronological order, the resulting behavioral motif may hold greater clinical significance. From these sequences we can characterize the frequency of transitions, or “grammar”, between behavioral modules in epileptic and non-epileptic mice.

We combined MoSeq with a novel workflow based on analyzing concatenated syllables sequences and network theory. This analysis reveals the emergence of an OCD like perseverative behavior in the kainic acid mouse model of epilepsy. We also find a hyperactivity phenotype associated with epilepsy specific modules of behavior that emerge and persist throughout chronic disease. Further, we quantified the emergence of epilepsy specific sedentary behaviors during disease progression. We also show that epilepsy leads to perturbed and increasingly fragile behavioral networks in chronic disease. Finally, we tested this pipeline by treating mice with the known anti-seizure medication carbamazepine (CBZ). Carbamazepine is a commonly prescribed anti-seizure medication used to treat not only epilepsy but also bipolar disorder and ADHD, disorders associated with increased rates of hyperactivity (26–28). We show that chronically epileptic mice treated with CBZ show reductions in racing behavior and partially rescued behavioral network dispersion. Together, this analysis provides a novel framework for analyzing MoSeq data which reveals clinically relevant behavioral phenotypes.

## Methods

### Systemic kainic acid model

FVB/Nhsd mice were weighed and singly housed in observation chambers for the duration of the injections. Mice were injected intraperitoneally (i.p.) with synthetic kainic acid (KA) (7.5mg/kg for FVB) dissolved in 0.9% saline (#7065, Tocris Bioscience, Bristol, United Kingdom). At twenty-minute intervals, mice were given 7.5 mg/kg injections of KA up to the third injection; the dosage was then reduced to 5.0mg/kg. Animals continued to receive 5.0 mg/kg KA every twenty minutes for up to 10 injections. If an animal experienced two or more Class V or VI seizures within a single twenty-minute cycle, the subsequent KA injection was skipped. Injections resumed for the next round unless the animal reached status epilepticus (SE), and animals were considered to have reached SE after experiencing at least five Class V or VI seizures within a 90-minute window. During induction of SE, seizures were scored using a modified Racine Scale (86) where I = freezing, behavioral arrest, staring spells; II = head nodding and facial twitches; III = forelimb clonus, whole-body jerks or twitches; IV = rearing; V = rearing and falling; VI = violent running or jumping behavior. KA mice were observed for 1-2 hours after SE was achieved. Animals were returned to their home cages post SE. In the days following injection, animals were weighed and injected with 0.9% saline (s.c.) if body weight decreased by more than 0.5g and given gel diet (cat #) to aid in recovery.

### Carbamazepine treatment of KA mice

Following status epilepticus, a cohort of mice were treated with carbamazepine (CBZ) for two weeks. These mice were recorded for MoSeq analysis prior and post CBZ treatment. CBZ (carbamazepine) (#4098, Tocris Bioscience, Bristol, United Kingdom) was dissolved in a solution of 30% PEG 400 (#39927-08-7, SIGMA, Darmstadt, Germany) in saline. CBZ was administered via i.p. injections at a dose of 35 mg/kg twice daily 4 hours apart for two weeks in chronically epileptic mice, 12-14 weeks post SE.

### Behavioral Seizure recording

To verify that mice had developed epilepsy, all 12- and 20-weeks post-SE mice were video recorded for behavioral seizures. Male and female FVB mice were singly housed in observation chambers for the duration of recording with access to food and water, and mice that did not exhibit at least 1 seizure per week (0.2/day) during the baseline recording period (or who died during recording) were triaged from the analysis. Videos were then reviewed and scored by an experimenter using the modified Racine scale, and only class IV-VI were recorded.

### Behavioral data acquisition

Behavioral data acquisition was performed similar to previous descriptions(21,23,25). In brief, mice were placed in an open arena (17 inch diameter and 16 inch walls, 14317, United States Plastics Corp) and recorded immediately after. The opaque enclosure was sandpapered (4664A16, McMaster-Carr) and spray painted black (54TG81, Krylon Industrial) to avoid image artifacts. The enclosure was cleaned between mice with 70% ethanol. Each animal was allowed to freely explore the enclosure for 20 min. For disease epilepsy progression experiments, the mice were recorded at the same time every day during the light cycle at 1.5-, 12-, and 20 after the inductions of status epilepticus. The recordings occurred during the interictal period, and no mice were observed to have behavioral seizures prior to placement in the recording enclosure.

### Behavioral recording

Data acquisition was performed as previously described(21,23,25). Briefly, each mouse was tracked in 3D using a Kinect 2 camera suspended .65 cm over the arena providing a top-down view. Raw data from the camera was then sent to the acquisition and analysis computer (Ryzen 7 5800X AMD 8-core with 16GB DDR4 RAM and EVGA GeForce RTX 3070 Ti FTW3 8GB graphics card) via USB 3.0 cables and depth frames were retrieved at 30 frames per sec.

### Motion Sequencing (MoSeq) Data extraction

The data preprocessing, extraction and modeling pipeline was performed as previously described(21,23,25). In brief, software was used to track the mouse in the enclosure and extract is position, orientation, height, size, and shape from the depth data. The 3D image of the mouse was extracted from that data using a previously described pipeline(21,23).

### MoSeq Data analysis – Transition Matrices

For each mouse we recorded the frequency with which one syllable transitioned into the next: thre frequency of “incoming syllable’ -> ‘outgoing syllable’. This data was used to generate a n x n transition matrix where n = number syllables which explain 99% of behavioral variance (see code).

### Network analysis – transition matrices

Individual non-normalized transition matrices were labeled to have “incoming_syllable number” along the side, and “outgoing_syllable number” along the top. Our individual non-normalized transition matrices match the structure of the matrices generated by MoSeq. Prior to network analysis we transformed transition matrices into a long format, with one column reserved for incoming syllable, another column for outgoing syllable, and a third column for the transition frequency (i.e. the number of transitions for that incoming outgoing pair of syllables). This output was then used to generate directed networks in cytoscape for each individual naïve and epileptic mouse at each timepoint. In these networks the nodes represented the syllables, and the edge represented a transition from one syllable to another, weighted by frequency. We also generated the same networks using the Python networkx library(29). The following network measures were collected: edge count, average neighbor degree, average degree connectivity, closeness vitality, betweenness centrality, and closeness centrality. Each metric was analyzed for significance using a Mann Whittney test. For Edge count at 12 weeks post SE, we also compared the first quartile and robust max (defined as the top three values).

### Edge count

Edge count is the total number of edges for each node. We calculated edge count in two different ways. For figure 6 and supplemental we used weighted edges, each edge representing a transition with the weight associated with the transition frequency in numbers. For figure 3, we replicated the edges for each node pairs by the number of transitions that edge pair had (for example, instead of incoming 2 outgoing 4 having one edge with a weight of 20, there would now be 20 edges all with a weight of 1).

### Average neighbor degree

Average neighbor degree is the average weighted edge count (*s_i_*) for the neighbors of a given. Since we used directed networks, the neighborhood (*N*_(*i*)_) of a node is determined based on incoming and outgoing nodes (*j*) of a given node (*i*), considering the weighted edge (*w*_j_) that links nodes *i* and *j* times the edge count (*k_j_*) of nodes *j*. This can be calculated for weighted networks using the following formula:

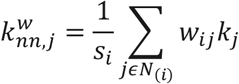

Average neighbor degree measures the connectivity of a network by measuring the edge count of the neighbors of a given node, higher average neighbor degree indicates increased network connectivity and reduced node dispersion. Since average neighbor degree does not consider node number for different networks, it cannot be compared across timepoints that have a different max number of nodes for their respective networks.

### Average degree connectivity

Average degree connectivity is average weighted edge count of all neighboring nodes (*N*_(*i*)_) surrounding a node (*i*) of a given weighted edge count (*s_i_*). The weighted edge (*w_ij_*) that links the node of interest (*i*) to its neighboring nodes (*j*) is multiplied by the total edge count (*k_j_*). Average degree connectivity can be calculated using the following formula:

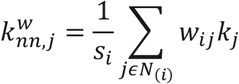

Average degree connectivity measures the connectivity of a network via the connectivity of nodes with a given edge count, networks with higher average degree connectivity are less dispersed and more connected.

### Closeness vitality

The closeness vitality of a node is the change in the sum of distances between all nodes when said node is removed from the network. This can be determined by calculating the wiener index of the graph with the node and without the node. The formula for the wiener index is below:

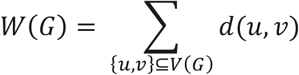

Wherein *G* represents the network, *V*(*G*) is the set of vertices in a graph, or nodes, and *d*(*u*, *v*) is the distance of the shortest path connecting the nodes. The removal of some nodes from the network leads to a disconnected network. When this occurs the closeness vitality of this node is coded as negative infinity. We accounted for these nodes by assigning all of them a large negative value, −1E99, during our analysis. By measuring closeness vitality, we can assess how vulnerable a network is to perturbations.

### Betweenness Centrality

Betweenness centrality measures the dependance of the network on a given node (*v*) by summing the fraction of node pairs shortest paths (*s*, *t*) that go through said node. The formula is displayed below:

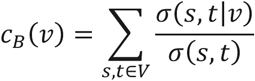

Were *v* is a set of nodes and *σ*(*s*, *t*) is the number of shortest paths, *σ*(*s*, *t*|*v*) is the number of shortest paths passing through the node (*v*) of interest. *V* cannot be *s* or *t*. By measuring betweenness centrality we can assess the importance of a given node to information flow in a network and wholistically the extent to which hub nodes exist within a network. Since this formula accounts for the total number of nodes in a network, it can be compared across different timepoints in epilepsy progression which also have different max node numbers for their networks.

### Closeness Centrality

The closeness centrality of a given node in a directed graph represents the independent centrality of a given node, by calculating the reciprocal of the average shortest incoming path to node (*u*) oval all (*n* − 1) reachable nodes. The formula for closeness centrality is listed below:

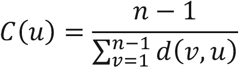

Wherein *d*(*v*, *u*) is the shortest path distance between nodes *v* and *u*. Closeness centrality allows us to measure how dispersed the nodes of a network are separate from edge counts.

#### Word clouds

Word clouds were generated using wordclouds.com. The size of the word was based on the number of times a given behavior was repeated in the syllable list. For instance, if ‘edge racing’ was repeated twice it would have a size of 2. See supplemental data for the syllable lists that generated the word clouds.

#### Syllable sequences

The sequence of all syllables that a mouse moves through in the recording period underlies the generation of transition matrices. We extracted syllable sequences for each mouse and recorded bouts of repetitive behavior.

Following syllable extraction from MoSeq, we quantified repetitive versus non-repetitive behavioral alternations using a sliding three-syllable window. This process was based on the method used to quantify spontaneous alternations in the spontaneous alternation Y maze test previously described(30), now applied to syllable space. Let the MoSeq-derived syllable sequence for a given animal be:

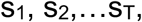

where each s_t_ denotes the integer-encoded syllable at time t. For each overlapping triplet

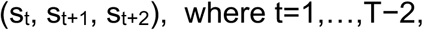

we classified the pattern as non-repetitive if the first and third syllables differed, and repetitive if they were identical. We formalized this using a binary indicator variable

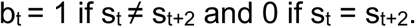

Thus, an example non-repetitive alternation such as 5,6,9 yields b_t_ = 1, whereas a repetitive alternation such as 12,1,12 yields b_t_ = 0. This procedure generates a binary sequence b_1_,…,b_T−2_ summarizing the alternation structure of the entire behavioral session.

To quantify overall behavioral flexibility, we calculated the fraction of non-repetitive alternations as:

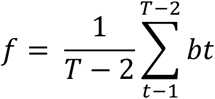

To assess the persistence of repetitive or flexible patterns, we next quantified the lengths of consecutive non-repetitive versus repetitive alternation bouts. Within the binary sequence b_t_, we defined a run as any maximal contiguous block of constant value (all 1s or all 0s). If a run begins at index s_k_ and ends at s_k+1_-1, its length is:

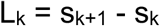

These run lengths were used to compute summary statistics for each animal and condition.

### Statistical analysis

For all statistical analysis—unless otherwise specified—*q* < 0.05 or *P* (corrected for multiple comparisons) <0.05 was considered statistically significant (\**P*/*q* < 0.05, \*\**P*/*q* < 0.01, \*\*\**P*/*q* < 10^−3^, and \*\*\*\**P*/*q* < 10^−4^). Data were graphed as means ± SD in Prism 10 software (GraphPad Software, La Jolla, CA). Statistical tests were also performed in Prism with the exception of the PERMANOVA test which was performed using the vegan package in R. Syllable usage, sequency analysis, and quartile analysis were analyzed by *t* test, two-way ANOVA/mixed-effects models, or one-way ANOVA, with the two-stage step-up method of Benjamini, Krieger, and Yekutieli (abbreviated BKY in figure legends). Kruskal-Wallis tests with Benjamini, Krieger, Yekutieli correction for data with >2 conditions. Analysis of length of consecutive repeats and non-repeats as well as network metric analysis was conducted using a PERMANOVA (999 permutations) using Kolmogorov-Smirnov distance with Benjamini-Hochberg correction for data with > 2 conditions. Fisher’s exact test was used for measuring the participation in behavioral tests with p-value reported. All statistical tests were two-tailed.

## Results

### Epileptic mice exhibit more repetitive alternations than naïve mice when navigating syllable space

Previous work using MoSeq in epilepsy has focused on changes in syllable usage between epileptic, naïve and drug treated mice(25). Though each syllable on its own has little clinical relevance, we reasoned that when sequenced together, concatenated syllables as behavioral motifs may hold greater significance. Therefore, for each sequence we defined non-repetitive behaviors as “non-repetitive alternations”, such that for a behavioral motif consisting of 3 sequential syllables, the first and third syllable are not the same (e.g. “rear down”->”scrunch”->”short lunge”). A motif where the first and third syllable are the same was classified as a “repetitive alternation” (e.g. “rear down”->“scrunch”->”rear down”) **(Fig 1A)**.

**Figure. 1.**
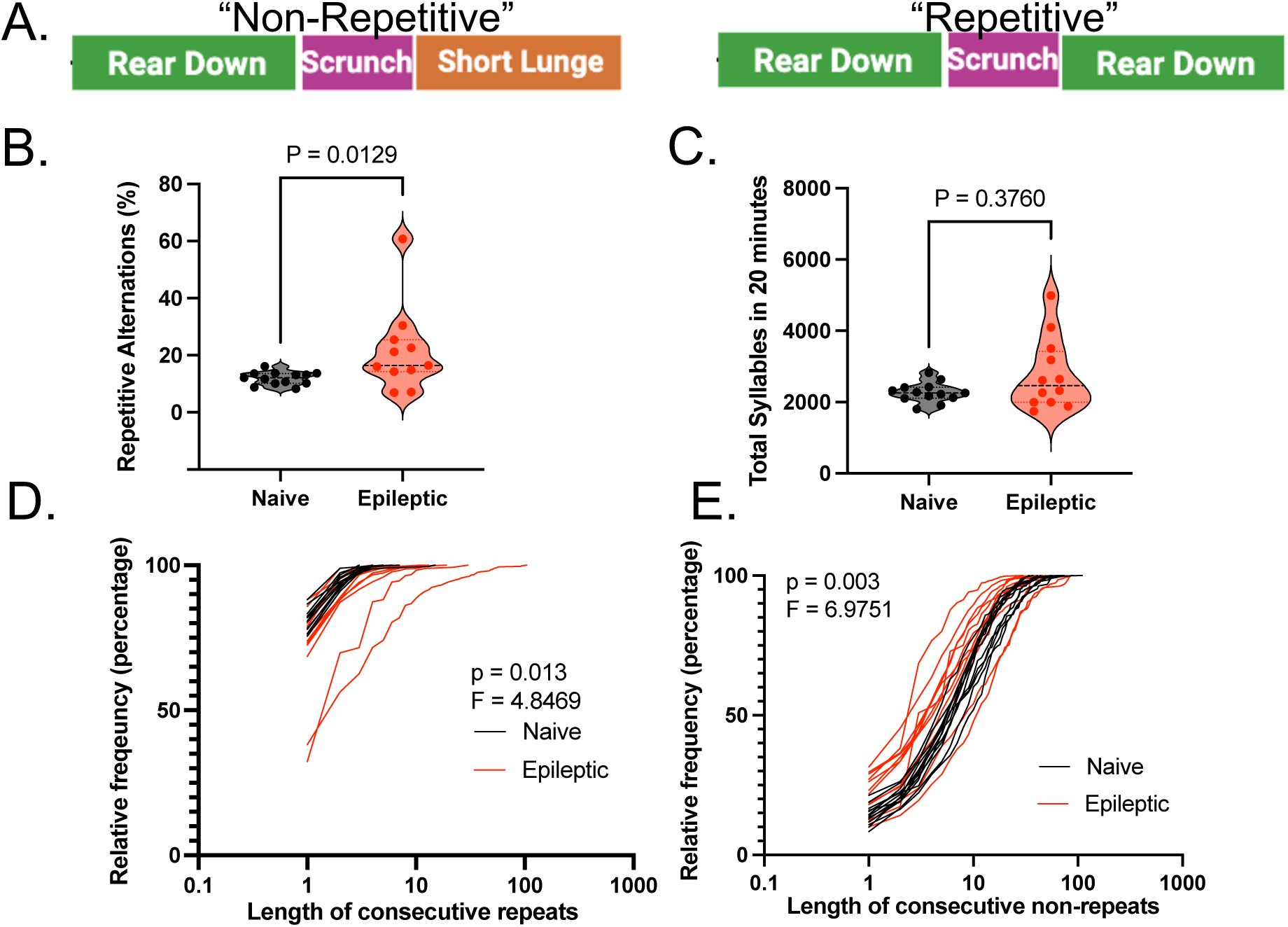
Epileptic mice have significantly more repetitive alternations than naïve mice. **(A)** Examples of non-repetitive or repetitive alteration using a three-syllable window**. (B)** Violin plots represent the percent repetitive alternations for naïve and epileptic mice (p = 0.0129) and **(C)** total sequence length (p = 0.3760). Each point represents a mouse and statistics performed using the Mann-Whitney test. **(D)** Cumulative of the length of consecutive repetitive alternations for epileptic and naive (Pseudo-F = 4.8469, p = 0.013) mice are depicted with each line representing a mouse. Statistics were performed using the PERMANOVA (999 permutations) based on the Kolmogorov-Smirnov distance. **(E)** Cumulative of the length of consecutive non-repetitive alternations for epileptic and naive (Pseudo-F = 6.9751, p = 0.003) mice are depicted with each line representing a mouse. Statistics were performed using the PERMANOVA (999 permutations) based on the Kolmogorov-Smirnov distance.

We find that epileptic mice have a significantly higher percent of ‘repetitive alternations’ compare to naïve mice (Mann-Whitney test p = 0.0129) **(Fig 1B)**. This was not the result of differences in total movement between the two groups (Mann-Whitney test, p = 0.3760) **(Fig 1C)**.

We then tested whether the increase in repetitive alternations was the result of extended bouts of repetitive behavior, also known as perseveration. Indeed, epileptic mice have significantly longer strings of consecutive repeats compared to age-matched naïve mice (PERMANOVA, p = 0.00.13, Pseudo-F = 4.8469) **(Fig 1D) (Fig S1)**. Therefore, epileptic mice are not only repeating behaviors more often, but they are also repeating behaviors for longer stretches of time. Epileptic mice also have significantly shorter strings of consecutive non-repetitive alternations compared to age-matched naïve mice (PERMANOVA, p = 0.003, Pseudo-F = 6.9751) **(Fig 1E) (Fig S1)**. This suggest that the ability of epileptic mice to have non-repetitive alternations is frequently interrupted by a repetitious alternation.

### Epileptic mice exhibit altered transition networks that prioritize a few nodes with high connectivity

We projected the sequential transitions from one syllable to the next as a directed and weighted network. Each node represents a syllable, and each edge represents a transition from one syllable to the next weighted by the frequency of that transition. Representative naïve and epileptic networks are shown in (**Fig 2A).** The representative epileptic network shows many low-weight edges and a few high weight edges whereas the naïve network shows a range of medium weight edges indicating that some epileptic mice may have nodes with abnormally strong connectivity (see supplemental figures for all individual epileptic and naïve networks). One way to analyze network connectivity is using edge count, a metric of how connected a given node is to its neighbors. For each individual mouse, we generated a cumulative frequency distribution of individual node edge counts **(Fig 2B)**. Analysis reveals that in the first quartile (representing nodes with lower edge counts) epileptic mice have significantly lower edge counts (Mann-Whitney Test, p = 0.0067) but at approximately 97^th^ percentile(i.e the robust max - representing nodes with high connectivity) epileptic mice have significantly higher edge counts than naïve mice (Mann-Whitney Test, p = 0.0126) **(Fig 2B)(Fig S2)**. This suggests that epileptic mice have sacrificed the integration of low connectivity nodes in favor of a few highly connected nodes.

**Figure 2.**
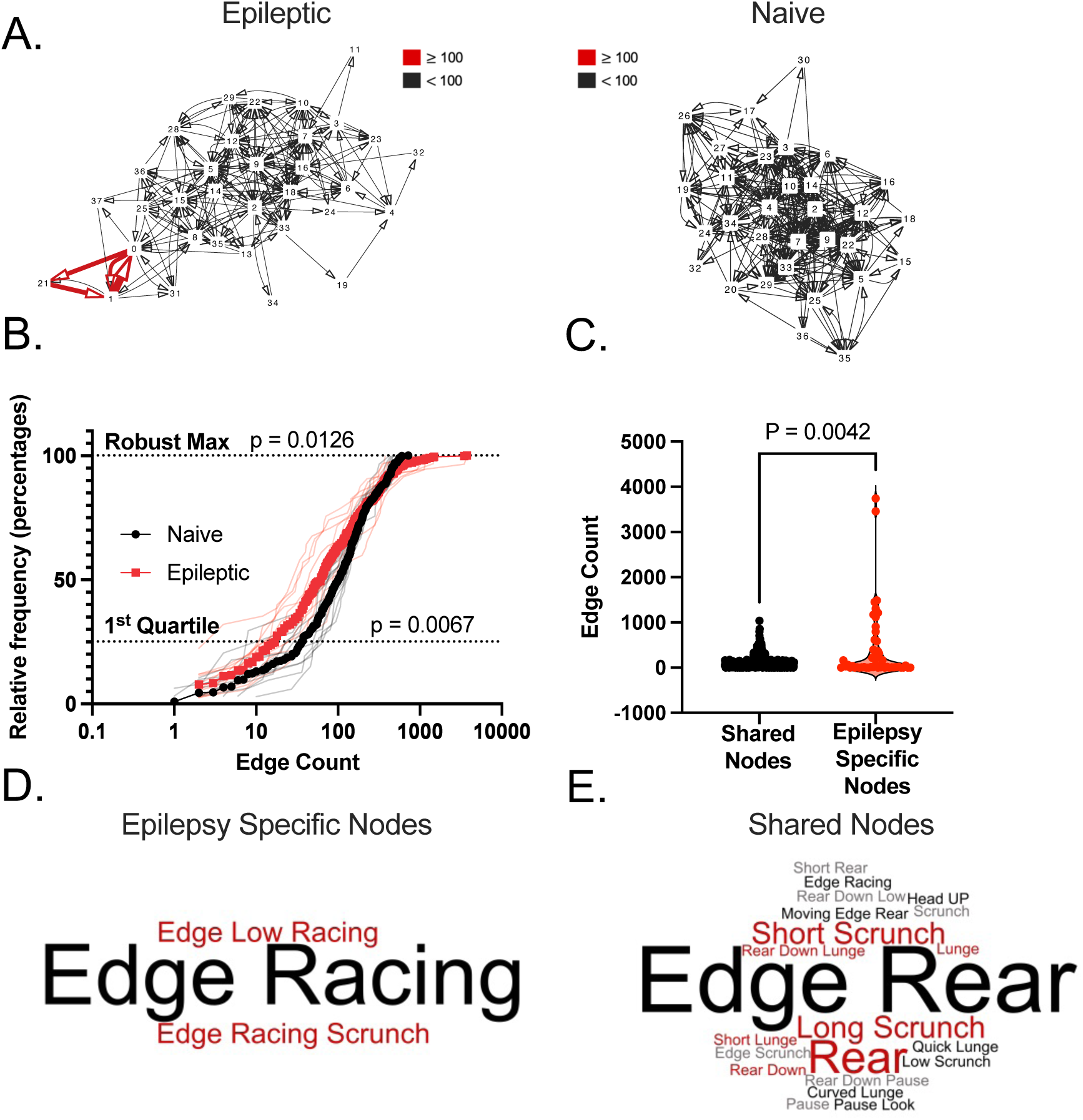
Epileptic mice exhibit abnormally connected networks with a racing phenotype. **(A)** Directed network diagrams of a single epileptic and naïve mouse where each node represents a ‘syllable’, and each edge represents a transition. Multiple transitions between nodes result in thicker and darker colored edges. **(B)** Cumulative of individual node edge counts for each individual epileptic and naïve mouse with the average distributions in bold and the individual distributions in the background. Statistical analysis for the 1^st^ Quartile (p = 0.0067) and Robust Max (top 3 values) (p = 0.0126) are depicted and were calculated using the Mann-Whitney test. **(C)** A violin plot represents the edge counts of nodes only present in epileptic mice compared to nodes shared between epileptic and naïve mice (p = 0.0042). Statistics calculated using Kolmogorov-Smirnov tests. **(D)** Word clouds created for the epilepsy specific nodes and the **(E)** nodes shared between epileptic and naïve mice. In both word clouds size indicates how often a node name is repeated in the node name list for a given group.

To determine whether the aberrant connectivity of chronically epileptic mice was the result of new behavioral motifs or the increased usage of behavioral motifs found in naïve mice we analyzed edge count of the 7 epilepsy specific nodes compared to the 30 nodes shared between naïve and epileptic mice. The epilepsy specific nodes have significantly higher edge counts than the nodes shared with naïve mice (K-S Test, p = 0.0042) **(Fig 2E).** The behavior these nodes represent are related to ‘edge racing’ **(Fig 2F)** and racing more broadly which was not represented in the rest of the naïve shared syllables **(Fig 2G).** This indicates that the aberrant connectivity of the epileptic networks could be due to the emergence of a racing phenotype in epileptic mice.

### ‘Racing’ syllables are upregulated in epileptic mice 12 weeks post SE and persists as epilepsy progresses

The presence of ‘racing’ syllables in epileptic networks lead the investigation of racing behavior more broadly. We classified all syllables such that any syllable labeled with a term that invokes rapid movement is called ‘racing’. To investigate whether racing syllable usage is prognostic of epilepsy we analyzed a 1.5-week post SE dataset for racing behavior prior to the emergence of chronic disease. We found racing is present in equal usage in both naïve and epileptic mice 1.5-weeks post SE (2-way ANOVA, q = 0.3531) **(Fig 3A).** In this 1.5-week post SE dataset all syllables were shared between epileptic and naïve mice indicating less of a divergence in behavioral phenotypes **(Fig S3**). At 12-weeks post SE this racing phenotype significantly decreases in naïve mice (2-way ANOVA, q = 0.0315) but continues to increase in epileptic mice (2-way ANOVA, q = 0.0156) **(Fig S3).** Racing syllable usage remains elevated 20-weeks post SE (2-way ANOVA, q = 0.0156).

**Figure 3.**
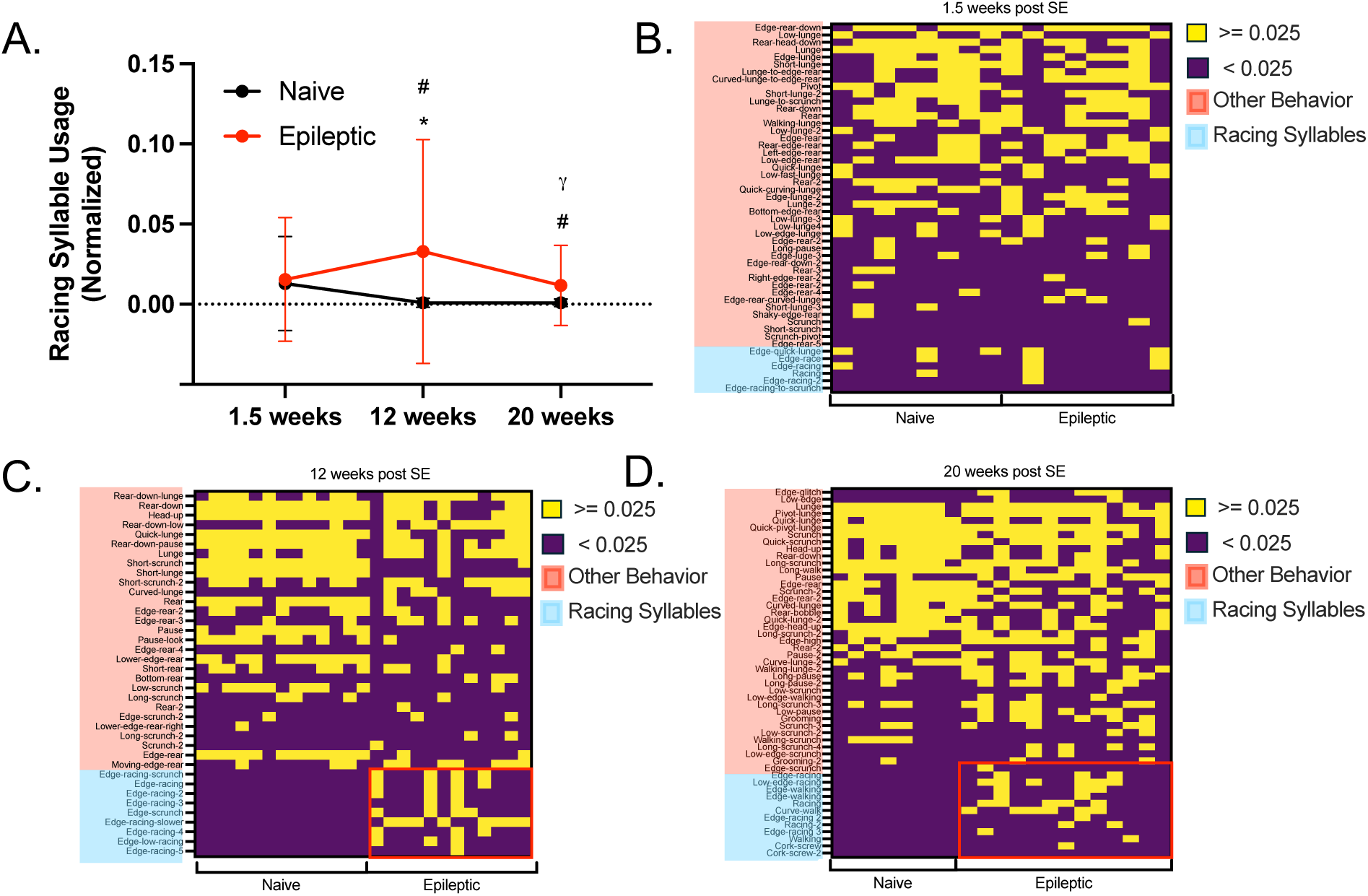
Racing syllables are upregulated in epileptic mice during chronic disease. **(A)** A summary line graph represents the racing syllable usage for naïve and epileptic mice 1.5-, 12-, and 20-weeks post status epilepticus. # indicates a significant difference between naïve and epileptic (q < 0.0001 at 12 weeks post SE, q = 0.0156 at 20 weeks post SE), * indicates a significant difference in epileptic racing syllable usage compared to 1.5 weeks post SE (q = 0.0067 at 12 weeks post SE). ψ indicates a significant difference between epileptic mice 12 weeks post SE and mice 20 weeks post SE (q < 0.0001). **(B-D)** Binarized heatmaps for racing syllable usage were generated for each timepoint with each column representing an individual mouse. The threshold of 0.025 normalized racing syllable usage determined by the highest racing syllable usage of any 12 weeks post SE naïve mice. The red outlined box shows the area for racing syllable for epileptic mice 12- and 20-weeks post SE. All syllable usage values were normalized to the total syllable usage for each mouse. All statistics were performed using a two-way ANOVA test with BKY post hoc.

To view racing behavior at a more granular level we generated a binarized heat map of racing and non-racing syllable usage **(Fig 3B-D).** Naive mice 12 weeks post SE, have the lowest racing syllable usage; therefore, we set the threshold for binarization at the highest normalized racing syllable for aged naïve mice 12 weeks post SE (binarizing threshold = 0.025). At 1.5-weeks post SE we see no difference in the binarized matrix between naïve and epileptic mice **(Fig 3B)**. However, at 12-weeks post SE every epileptic mouse shows usage of at least one racing syllable whereas no naïve mice show racing. Further, those epileptic mice 12 weeks post SE that use multiple racing syllables show a clear deficit in non-racing syllable usage (Mann-Whitney Test, p = 0.0061) **(Fig S4).** Once the mice have progressed to 20-weeks post SE, all but 2 epileptic mice still have 1 or more racing syllables **(Fig 3D)**.

### Mice shift to a more sedentary phenotype as epilepsy progresses from 12 to 20 weeks post SE

We find that as epilepsy progresses mice become increasingly sedentary and at 36-weeks post SE, they are unable to participate in traditional Y maze tests **(Table S1)**. To study this further, we generated a directed network for mice 20 weeks post SE. Once again there are a selection of 6 nodes that only appear in the epileptic networks. These epilepsy specific nodes have lower edge counts compared to nodes shared with naïve mice (K-S Test, p < 0.0001) **(Fig 4A)**. This contrasts with 12-weeks post SE when epilepsy specific nodes were aberrantly connected. Whereas epilepsy-specific nodes at 20-wks post SE contained ‘racing’ terms as seen at 12-weeks, the 20-weeks nodes also contained terms related to ‘scrunching’ **(Fig 4B)**. Shared nodes show a range of behaviors including ‘edge-rear’ and ‘long-scrunch’ **(Fig 4C)**. The prominence of the ‘scrunch’ behavior in the epilepsy specific nodes 20-weeks post SE links disease and sedentary behaviors in a way not seen 12-weeks post SE. To further investigate the differences between epileptic mice 12- and 20-weeks post SE we analyzed the total sequence length for epileptic mice at each time point. We show that 20-week post SE epileptic mice have significantly shorter sequence lengths than mice 12-weeks post SE (Mann-Whitney Test, p = 0.0205) **(Fig 4D)**. To test that this was due to disease and not aging per se we compared total sequence length between age match naïve mice at each time point and found no difference in total sequence length (Mann-Whitney Test, p > 0.9999) **(Fig 4E)**. This indicates that 20 weeks post SE epileptic mice move through less total syllable space than 12-week post SE mice, potentially due to a sedentary effect, despite the presence of racing syllables.

**Figure 4.**
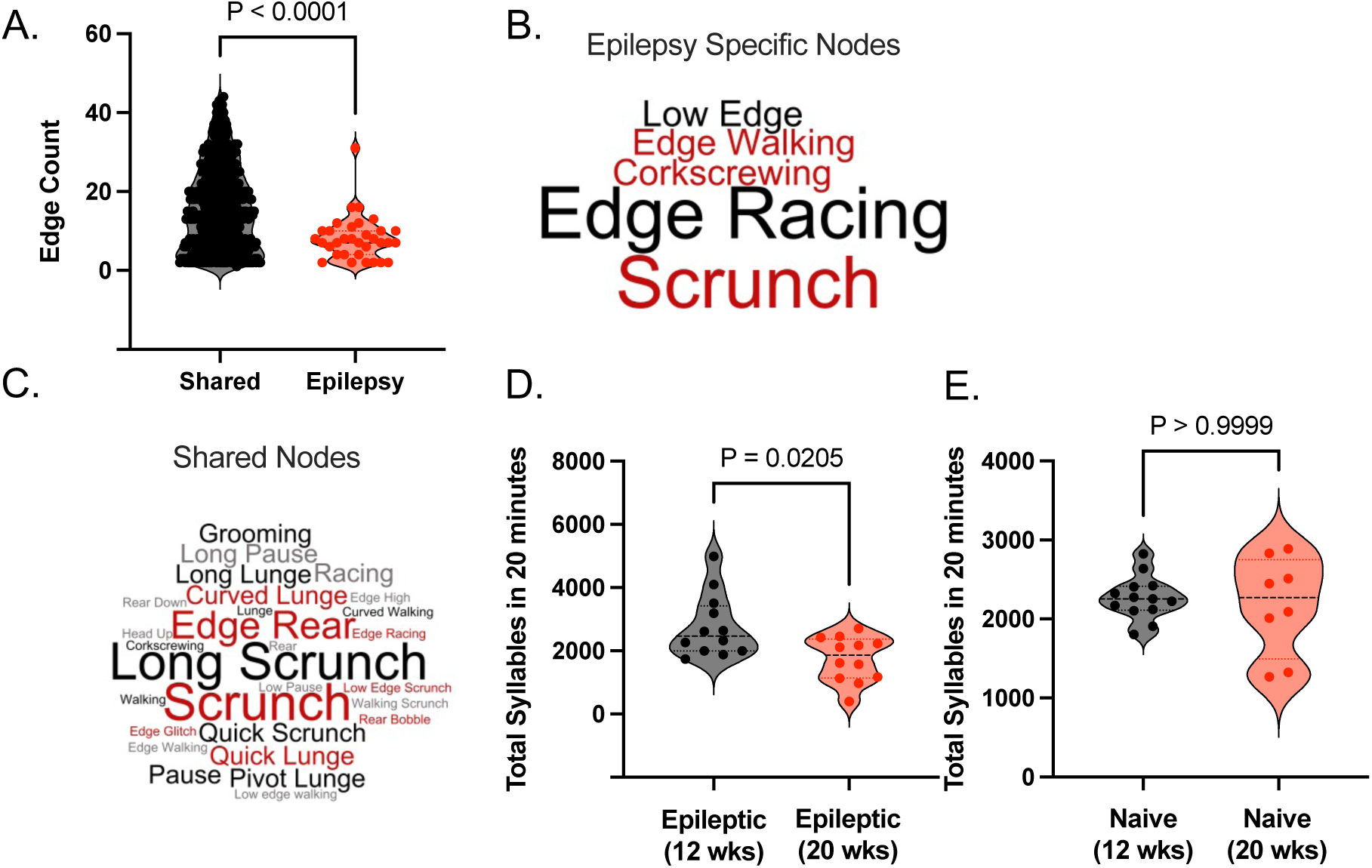
As epilepsy progresses to 20 weeks post SE syllables shift to a more sedentary profile. **(A)** Violine plots shows the reduction in edge count for epilepsy specific nodes at 20 weeks post status epilepticus (p < 0.0001). Each point represents an epilepsy specific nodes’ edge count. Depictions of word clouds showing the behaviors in **(B)** epilepsy specific nodes and **(C)** shared nodes for 20 weeks post SE. **(D)** Violin plots shows that sequence lengths are shorter from 12 to 20 weeks post status epilepticus (p = 0.0205) **(E)** while naïve mouse lengths are statistically the same for each timepoint (p > 0.9999). Each point represents a mouse. All statistics done using a Kolmogorov-Smirnov test.

### Network analysis reveals increased fragility of behavioral networks as epilepsy progresses

To further interrogate the network architecture of behavior through the course of epilepsy, we measured degree, average degree connectivity, and average neighbor degree at 1.5, 12 and 20 weeks post SE **(Fig S5) (Fig S6)**. We find a reduction in degree at 12 weeks post SE (PERMANOVA, p = 0.004, Pseudo-F = 5.615) **(Fig S5) (Fig S6)** matching our quartile analysis **(Fig 2B)**. However, closeness vitality, closeness centrality, and betweenness centrality stood out as significantly perturbed measures across disease progression **(Fig S7)**. Closeness vitality is the change in the sum of shortest distances between all node pairs when excluding that node and is a measure of network fragility i.e. how easily a network is perturbed when a node is removed. There was no difference in closeness vitality between naïve and epileptic networks at 1.5 weeks post SE **(Fig 5A)** (PERMAVOA, p = 0.384, Pseudo-F = 1.0129). However, we see an increase in epileptic closeness vitality at 12 weeks post SE (PERMAVOA, p = 0.001, Pseudo-F = 10.37) **(Fig 5B)**. This difference extends to 20-weeks post SE (PERMANOVA, p = 0.021, Pseudo-F = 5.4022) (**Fig 5C**). Additionally at 20-weeks post SE epileptic mice had a greater proportion of nodes that when removed resulted a disconnected network compared to naïve mice (Mann-Whitney, p = 0.0095) **(Fig S8)** suggesting that as epilepsy progresses the behavioral networks of mice are increasingly fragile.

**Figure 5.**
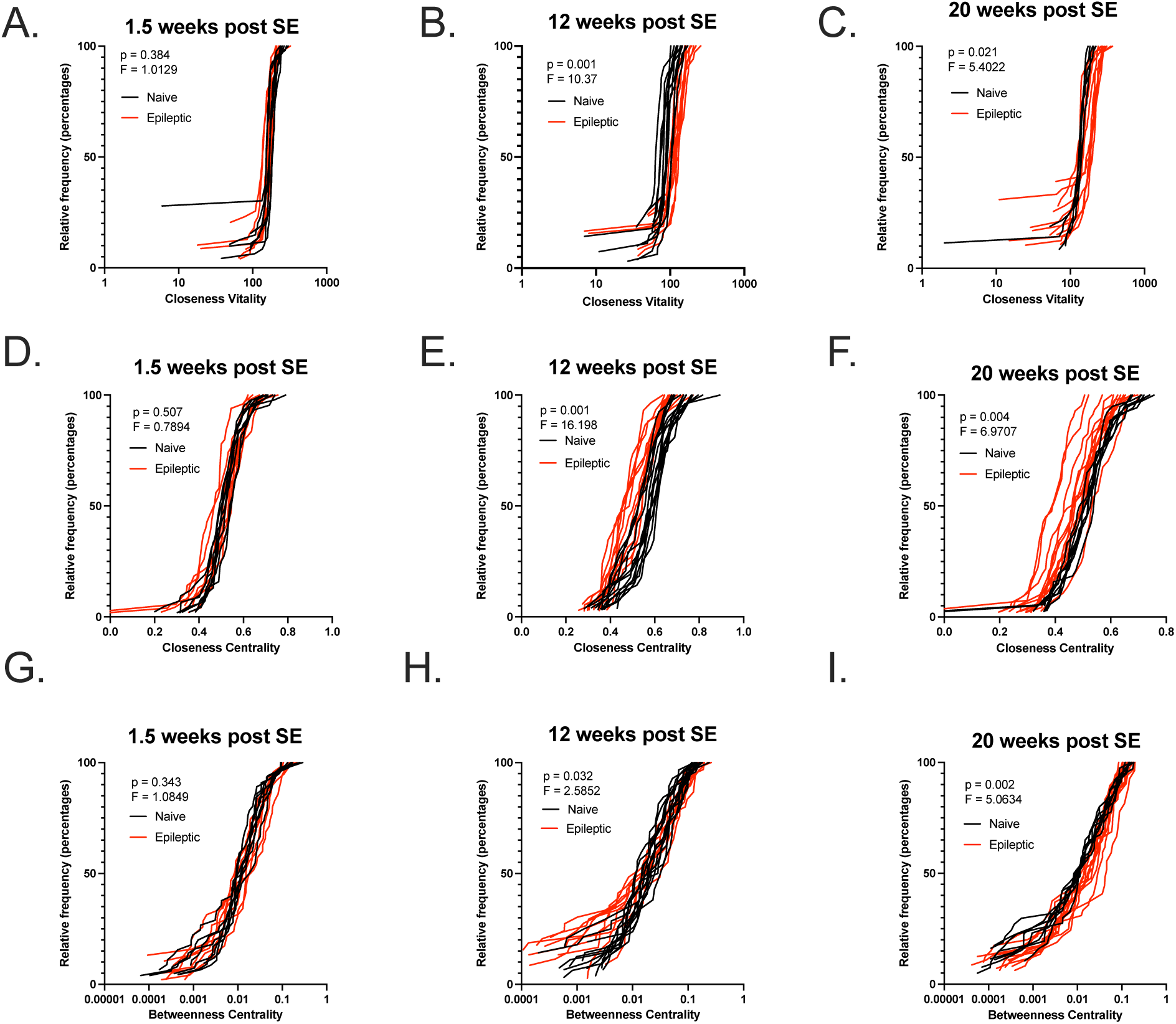
As epilepsy progresses behavioral network fragility increases. Network metrics for 3 different time points (1.5-, 12-, and 20-weeks post SE) with each line representing a mouse. Closeness vitality at **(A)** 1.5-weeks post SE (Pseudo-F = 1.0129, p = 0.384), **(B)** 12-weeks post SE (Pseudo-F = 10.37, p = 0.001), and **(C)** 20-weeks post SE (Pseudo-F = 5.4022, p = 0.021). Closeness centrality at **(D)** 1.5 weeks post SE (Pseudo-F = 0.7894, p = 0.507), **(E)** 12-weeks post SE (Pseudo-F = 16.198, p = 0.001), and **(F)** 20-weeks post SE (Pseudo-F = 6.9706, p = 0.004). Betweenness centrality at **(G)** 1.5-weeks post SE (Pseudo-F = 1.0849, p = 0.343), **(H)** 12-weeks post SE (Pseudo-F = 2.5852, p = 0.032), and **(I)** 20-weeks post SE (Pseudo-F = 5.0634, p = 0.002). All statistics performed using the PERMANOVA (999 permutations) based on the Kolmogorov-Smirnov distance.

We measured closeness centrality for each node of individual epileptic and naïve mice for each timepoint during disease progression plotting individual values as well as the average per condition (1.5-, 12-, and 20-weeks post SE) **(Fig 5D-F)**. Closeness centrality refers to the reciprocal of the average shortest paths over all reachable nodes. Thus, the higher the closeness centrality of a node, the shorter the path to reach any other node in the network and the more central is the node. We observe no difference in closeness centrality between naïve and epileptic mice at 1.5-weeks post SE (PERMANOVA, p = 0.507, Pseudo-F = 0.7894) **(Fig 5D)**. At 12-weeks post SE epileptic networks showed significantly decreased closeness centrality (PERMANOVA, p = 0.001, Pseudo-F = 16.198) **(Fig 5E)**. This trend extends to 20 weeks post SE (PERMANOVA, p = 0.004) **(Fig 5F)** indicating that the networks of epileptic mice are less centralized with fewer nodes close to each other.

We measured betweenness centrality for each node of individual epileptic and naïve mice for each timepoint during disease progression **(Fig 5D-F)**. Betweenness centrality refers to the proportion of all shortest paths in a network that go through a given node; networks with higher increased betweenness centrality have more nodes that act as necessary bridges or bottlenecks. While we observed no difference at 1.5 weeks post SE (PERMANOVA, p = 0.343, Pseudo-F = 1.0849) **(Fig 5D)**, at 12-weeks post SE epileptic networks displayed increased betweenness centrality (PERMANOVA, p = 0.032, Pseudo-F = 2.5852) **(Fig 5E)**. As epilepsy progresses to 20 weeks post SE epileptic mice have significantly increased betweenness centrality compared to age-matched naïve mice (PERMANOVA, p = 0.002, Pseudo-F = 5.0634) **(Fig 5F)** indicating that epileptic networks rely heavily on a selection of nodes for information flow, or movement, within the network.

### Treatment with Carbamazepine rescues epilepsy-associated hyperactivity and network fragility

To test whether the increased repetition, racing, and network fragility were the product of epilepsy associated hyperactivity we decided to treat our mice with the FDA approved anti-seizure medication carbamazepine (CBZ). Importantly, CBZ has also been used to treat a variety of psychiatric disorders including but not limited to, ADHD and mania in bipolar – both conditions associated with hyperactivity. Therefore, we hypothesized that as a stabilizing medication, CBZ may suppress some of the hyperexcitability phenotypes we observe in our epileptic mice. Analysis of syllable usage between naïve, epileptic and CBZ treated mice revealed that those syllables that CBZ rescued were mainly those upregulated during epilepsy and predominately include behaviors such as “racing” and “scrunch” **(Fig 6A) (Fig S9)** Conversely the syllables that failed to be rescued by CBZ treatment were mainly those that were downregulated during epilepsy and include behaviors such as “rear” and “lunge” **(Fig 6B) (Fig S10)**. This indicates that while CBZ is effective at suppressing behaviors upregulated by epilepsy it fails to restore behaviors that have been lost during disease. We also analyzed whether CBZ was effective at supressing the upregulation of “racing” behavior in chronic epilepsy and show that CBZ treatment significantly reduces normalized racing syllable usage compared to epileptic mice (Kruskal-Wallis, q = 0.0024) **(Fig 6C).** This shows that treatment with CBZ suppresses racing behavior overall. In addition, we found few off-target syllable effects in our analysis **(Fig S11) which we defined as** We analyzed the effect of CBZ on repetitive behaviors and found no significant effect outside of a reduction in overall number of syllables **(Fig S12) (Fig S13)**.

**Figure 6.**
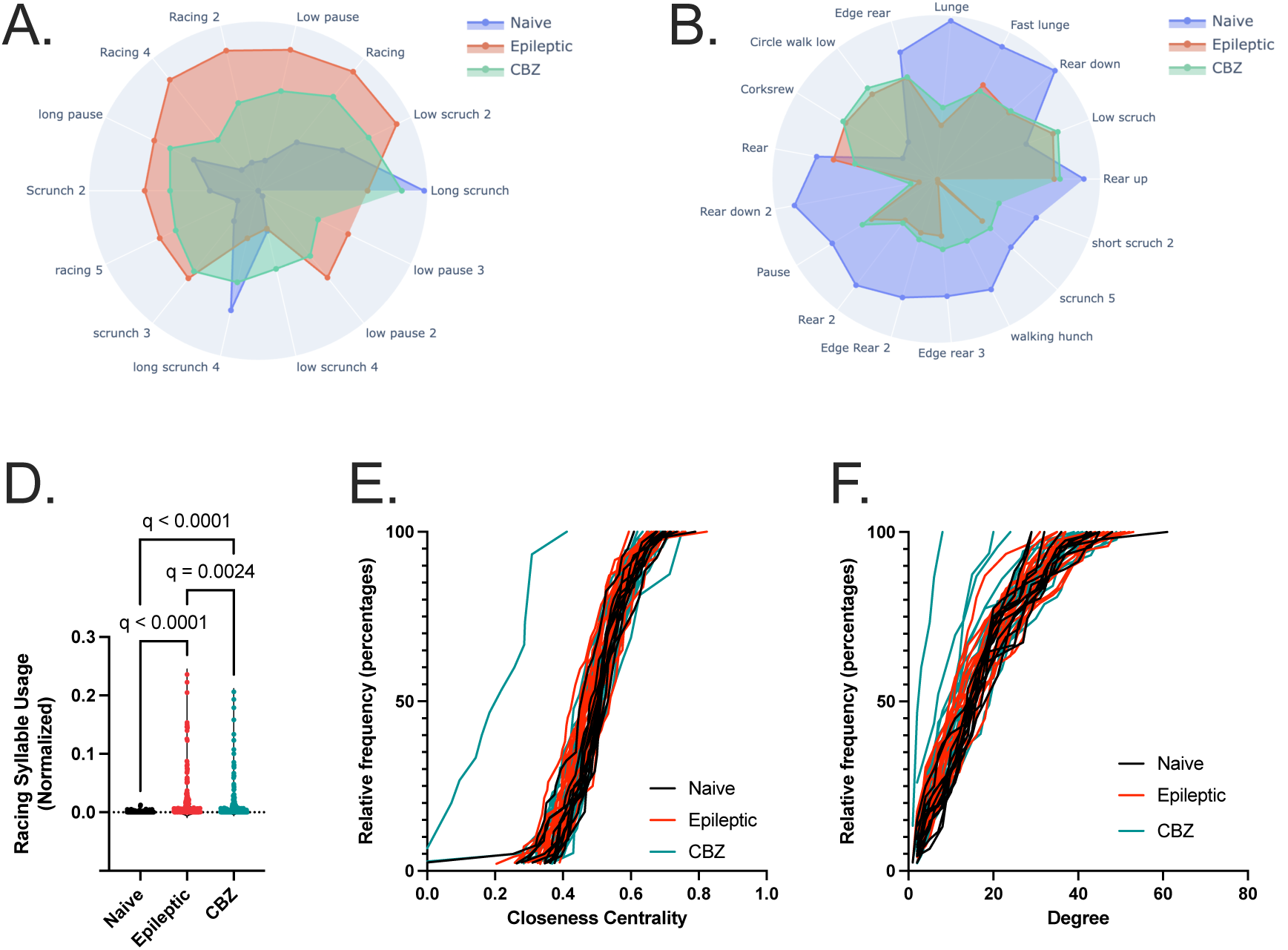
CBZ rescues behavioral alterations associated with hyperactivity. Polar plots depict the syllable **(A)** rescued and not rescued **(B)** by CBZ treatment in chronic epilepsy. Distance of each point from the center of the polar plot to the edge represents syllable usage. **(C)** Bar graph shows the effect of CBZ of normalized racing syllables in chronically epileptic mice (naïve and epileptic q < 0.0001) (naïve and CBZ, q < 0.0001) (epileptic and CBZ, q = 0.0024). Statistics performed using the Kruskal-Wallis Test with BKY post-hoc. Each dot represents a syllable. Network metrics for 3 different conditions (naïve, epileptic, and CBZ treated) with each line representing a mouse. **(D)** Closeness centrality (naïve and epileptic, Pseudo-F = 4.0488, p = 0.048) (naive and CBZ, Pseudo-F = 1.5752, p = 0.1995) (epileptic and CBZ, Pseudo-F = 0.722, p = 0.6969). **(E)** Degree (naïve and epileptic, Pseudo-F = 4.7831, p = 0.012) (naive and CBZ, Pseudo-F = 4.9102, p = 0.012) (epileptic and CBZ, Pseudo-F = 1.0863, p = 0.374). Network metrics statistical analysis performed using the PERMANOVA (999 permutations) based on the Kolmogorov-Smirnov distance with a Benjamini-Hochberg post hoc.

We analyzed average degree connectivity, average neighbor degree, betweenness centrality, closeness vitality **(Fig S14) (Fig S15)**, closeness centrality and degree of directed networks for our naïve, epileptic, and CBZ treated mice **(Fig 6D-F)**. We found that once again epileptic mice have significantly lower closeness centrality than age matched naïve mice (PERMANOVA, p = 0.048, Pseudo-F = 4.0488) **(Fig 6D).** This significant difference is no longer present upon treatment with CBZ (naïve and CBZ, PERMANOVA, p = 0.1995, Pseudo-F = 1.5752) **(Fig 6D)**, however CBZ treated mice are also statistically indistinguishable from epileptic mice, potentially indicating a partial rescue (epileptic and CBZ, PERMANOVA, p = 0.722, Pseudo-F = 0.6969). We do observe a reduction in the edge counts in the behavioral networks of epileptic mice compared to naïve mice (PERMANOVA, p = 0.012, Pseudo-F = 4.7831). Treatment with CBZ fails to rescue the reduction in degree for epileptic networks compared to naïve mice (PERMANOVA, p = 0.012) leaving CBZ treated mice indistinguishable from untreated epileptic mice (PERMANOVA, p = 0.374, Pseudo-F = 1.0863) **(Fig 6E)**. This is in line with other data showing that CBZ treated mice move through less total syllables during the 20 min recording time **(Fig S12)**. There is no significant difference between naïve and epileptic mice for either closeness vitality or betweenness centrality (PERMANOVA, p = 0.6542 and p = 0.5085 respectively) **(Fig S14) (Fig S15).** We also found not difference in the number of nodes which upon removal resulted in a disconnected network during our closeness vitality analysis **(Fig S16).**

## Discussion

In this study we applied a novel network analysis approach to characterize mouse behavior in MoSeq analysis of disease progression in the systemic kainic acid mouse model of epilepsy. Our analysis reveals that chronically epileptic mice engage in perseverative behaviors and have altered behavioral networks characterized by the emergence of epilepsy specific ‘racing syllables’. We performed this analysis at three different timepoints in disease (1.5-, 12-, and 20-weeks post SE) to discover differences in epilepsy specific behavioral phenotypes and behavioral networks as disease progresses. We validated this pipeline by showing that treatment with CBZ reduces multiple measures of hyper-activity, consistent with CBZ’s uses in other contexts such as ADHD and bipolar disorder (26–28). A sequence and network-based approach to MoSeq data in a disease framework yield results with relevance to comorbidity analysis.

### Sequence analysis reveals perseverative behaviors 12 weeks post SE

Patient reports as well as clinical studies have shown a correlation between psychiatric comorbidities, patients’ overall stigmatization, and reduced quality of life (10–12,31,32). However, many behavioral assays for comorbidities, such as depression, anxiety, and compulsive behaviors, in rodent models can only interrogate limited outcomes. These approaches utilize discrete tests that measure the ability to perform a specific well-defined task, such as the sucrose preference test, elevated plus maze, and marble burying test. Prior work by the Soltesz lab has demonstrated the ability of MoSeq to differentiate between epileptic and naïve mice in both acquired and genetic mouse models of epilepsy based on syllable usage(25). We expand on this work by testing whether analysis of concatenated syllables for repetitive alternation can further differentiate naive and epileptic mice. We find that epileptic mice repeat behaviors more often and for longer consecutive stretches than age-matched naïve mice **(Fig 1B, D).** Others have shown that beyond syllable usage, analysis of transitions from one syllable to the next can be useful(21). We posit that this approach holds increased clinical relevance. A mouse ‘scrunch’ may not translate to a patient with epilepsy, but the emergence of behavioral motifs, such as repetition, may be better mapped on to the phenotypes of disease. For instance, patients with epilepsy have higher rates of automatisms, defined as brief, repetitive, unconscious behaviors(33,34). These automatisms are generally associated with ictal events in both temporal lobe epilepsy as well absence seizures(33,34). TLE patients also have increased rates of obsessive-compulsive disorder (OCD) (11-34.5%) compared to the general public (0-3%)(9,35). Both of these clinical readouts could be plausibly mapped onto the repetitive behavior observed in our MoSeq analysis (8).

### Epilepsy specific syllables perturb behavioral networks 12 weeks post SE

We show that epileptic mice have perturbed behavioral networks resulting from the emergence of epilepsy specific syllables. We find that epileptic behavioral networks, generated from the transitions between incoming and outgoing syllables, have a few nodes with edge counts much greater than those present in naïve mice **(Fig 2B)**. Thus, high edge counts often connect 2 or 3 nodes in the network suggesting that repetition may be driving this result. Furthermore, a subset of these highly connected nodes is specific to epileptic mice and not present in any naïve animals **(Fig 2C)**. This indicates that the perseverative phenotype is the result of novel syllables being introduced into the network. We find that epilepsy specific syllables are 1) hyper connected and 2) are defined by a ‘racing’ and an ‘edge racing’ phenotype: mice sprint around the edge of the arena. This aligns with reported observations of kindled mice which use novel syllables that are not present in non-kindled mice in MoSeq recordings(25): the racing syllables we report match the novel syllables of ‘wild running’ seen in mice kindled 5 minutes before MoSeq recording(25,36).

### Racing syllable usage decays with age in naïve mice but persist in disease

We find that racing syllable usage is not significantly different between epileptic mice early in disease and age-matched naïve animals. However, whereas naïve mice show an age dependent decrease in racing syllable usage, epileptic mice persist in racing behavior **(Fig 3A)**.

This suggests that racing syllables are a poor biomarker of epilepsy early in disease. MoSeq was previously used to analyze changes in syllable usage during the first 4 weeks of disease in the intra-hippocampal kainic acid model(25). We expand our analysis of disease progression in the systemic kainic acid model of epilepsy to 5 months. Furthermore, we noticed that mice manifesting multiple racing syllables showed a lower usage of non-racing syllables **(Fig S5).** It is tempting to hypothesize that the racing behavior may be connected to hyperactivity in epileptic animals. Others studying hyperactivity in rats and ADHD in humans have shown a link between interictal spiking and behavioral alterations (37–39). Levetiracetam, an FDA approved anti-seizure medication, has been shown to successfully suppress interictal spiking in 63% of patients with ADHD with and without concurrent seizures(40). Of the patients with suppressed interictal spiking 59% showed behavioral improvements(40). Future work will test the hypothesis that racing behavior is the result of interictal epileptiform discharges.

### Epilepsy progression from 12-to 20-weeks post SE is characterized by the emergence of a sedentary phenotype

We find that as mice progress from 12-to 20-weeks post SE, epilepsy specific syllables manifest a reduction in edge count, signaling less connectivity to the overall network **(Fig 4A)**. This change corresponds with the emergence of epilepsy specific scrunching behaviors alongside the preexisting racing syllables **(Fig 4B-C)**. We believe that this change is due to disease progression rather than aging per se because naïve mice maintain their total syllable sequence length from 12-20 weeks post SE while epileptic mice show a significant reduction **(Fig 4D-E)**. Conventional y-maze tests only reveal a sedentary transition at 36 weeks post SE **(Table S1)**. These data highlight the ability of MoSeq, with its focus on spontaneous behavior, to detect a shift in mouse behavior earlier in disease progression compared to conventional behavioral assays.

### Epileptic mice show increased behavioral network fragility

Epileptic networks have multiple properties that diverge from naïve networks, and these persist during disease progression. For instance, epileptic mice exhibit more ‘fragile’ behavioral networks: they are populated by nodes with increased closeness vitality. The closeness vitality of a node is the change in the sum of distances between all node pairs when excluding that node(41). At 1.5-weeks post SE we observe no difference in closeness vitality in epileptic mice compared to age-matched naïve mice, however, by 12-weeks post SE epileptic mice have significantly increased closeness vitality which persists to 20-weeks post SE **(Fig 5A-C)**. This suggests that as disease progresses epileptic behavioral networks become increasing fragile. This change is accompanied by an increase in the number of ‘privileged’ nodes which when removed result in a disconnected network at 20-weeks post SE (Fig S8) Thus, removal of a high closeness vitality syllable in an epileptic network result in cascading effects, including the increased dispersion of the network. Additionally, as disease progresses from 1.5 weeks post SE to 12- and 20-weeks post SE epileptic networks show a decrease in closeness centrality **(Fig 5D-F)** – a measure of how tightly the nodes of a network are connected (41). Networks with low closeness centrality nodes tend to be dispersed and loosely connected. Conversely, epileptic behavioral networks increased betweenness centrality at 12-, and 20 weeks post SE **(Fig 5G-I).** Betweenness centrality is a measure of the proportions of shortest paths which go through a given node, i.e. betweenness centrality can measure the number of bridges and bottles necks within a given network(41). While one may hypothesize that closeness centrality and betweenness centrality must be correlated this is not necessary for weighted networks. For instance, in our reference epileptic network we see a group of peripheral nodes (0, 21, and 1) that are connected by a few high weight edges with node 0 serving as a bridge between these nodes and the rest of the network **(Fig 2A).** The peripheral location of these bridge nodes can contribute to the overall fragility of the network, since if node 0 is removed the network is more prone to disconnection **(Fig 2A)**. Together, these results suggest that as disease progresses the network loses resilience, i.e. if a node is removed from the network that loss has a greater impact on the topology of the network.

### Treatment with carbamazepine rescues behavioral alterations associated with hyperactivity

Carbamazepine is an antiseizure medication that has been shown to reduce hyperactivity and mania in patients with ADHD and bipolar disorder respectively(26–28). We find that treatment with CBZ is most effective at supressing behaviors elevated during epilepsy but often fails to restore normal behaviors dampened during disease (**Fig 6A-B).** More specifically, CBZ is able to suppress the aberrant increase in racing syllable usage in epileptic mice **(Fig 6C)**. This is in line with CBZ’s known effects in a rodent model with hyperactivity (42) as well as its clinical applications outside to reduce hyperactivity in patients(26). We also show that treatment with CBZ partially rescues the closeness centrality of epileptic mice **(Fig 6D).** This indicates that CBZ’s stabilizing effects may lead to less dispersed behavioral networks. However, it must be noted that mice treated with CBZ continued to have increased repetition both in percentage and length of consecutive repeats **(Fig S10).** While there is rodent data indicating that CBZ reduces marble burying this has not translated to significant patient results(43,44). Furthermore, CBZ treated mice moved through fewer syllables in 20 minutes than naïve mice which is in line with the known CBZ side effect of sedation and ataxia **(Fig S9)**(26,45). This effect is mirrored by the reduction in degree, or edge count, seen in the behavioral networks CBZ treated mice compared to naïve mice **(Fig 6E)**. Since CBZ treated mice move through fewer total syllables it would follow that they also have fewer transitions in their associated behavioral networks.

While others have used MoSeq as a high throughput screening tool to distinguish between different drug treatments in both naïve and epileptic mice (23,25), to our knowledge, the studies herein are the first to1) test the behavioral effects of CBZ with MoSeq and 2) expand our analysis beyond on and off target effects on syllable usage.

In sum, we have tested the use of network and syllable sequence-based analysis during disease progression in a mouse model of acquired epilepsy. Our findings of 1) increased repetitive behaviors, 2) increased racing, and 3) perturbed behavioral networks in chronic epilepsy map on to clinical comorbidities present in epilepsy patients. Furthermore, we show that treatment with a known anti-seizure medication carbamazepine reduces hyperactivity in line with its stabilizing activity in both bipolar disorder and ADHD. We propose that this framework can be used in other mouse models of progressive neurological diseases to derive clinically relevant behavioral outputs.

## Notes

**Funding:** Supported by CURE (AR), Lily’s Fund (AR), NIH grant T32 GM14103 (JK), R21NS139176 (AR) 1R01NS108756 (AR), R01NS108756 and R01NS102937 (JM).

### Competing Interest Statement

The authors have declared no competing interest.

### Summary of Updates

abstract updated to clarify rational. no other revisions

## References

1. Fiest F, Sauro K, Weibe S, Patten S, Kwon CS, Pringsheim T, et al. Prevalence and incidence of epilepsy: a systematic review and meta-analysis of international studies. 2016. Report.

2. Maguire J. Mechanisms of Psychiatric Comorbidities in Epilepsy. In: Current Topics in Behavioral Neurosciences. Springer Science and Business Media Deutschland GmbH; 2022. p. 107–44. doi:10.1007/7854_2020_192 PubMed PMID: 33723802.

3. Weatherburn CJ, Heath CA, Mercer SW, Guthrie B. Physical and mental health comorbidities of epilepsy: Population-based cross-sectional analysis of 1.5 million people in Scotland. Seizure. 2017 Feb 1;45:125–31. doi:10.1016/j.seizure.2016.11.013 PubMed PMID: 28024199.

4. Mula M, Kanner AM, Jetté N, Sander JW. Psychiatric Comorbidities in People With Epilepsy. Neurol Clin Pract. 2021;11(2):E112–20. doi:10.1212/CPJ.0000000000000874

5. Kleen JK, Scott RC, Holmes GL, Lenck-Santini PP. Cognitive and behavioral comorbidities of epilepsy. Epilepsia. 2010;51(SUPPL. 5):79. doi:10.1111/j.1528-1167.2010.02865.x

6. Ottman R, Lipton RB, Ettinger AB, Cramer JA, Reed ML, Morrison A, et al. Comorbidities of epilepsy: Results from the Epilepsy Comorbidities and Health (EPIC) survey. Epilepsia. 2011;52(2):308–15. doi:10.1111/j.1528-1167.2010.02927.x PubMed PMID: 21269285.

7. Miller DJ, Komanapalli H, Dunn DW. Comorbidity of attention deficit hyperactivity disorder in a patient with epilepsy: Staring down the challenge of inattention versus nonconvulsive seizures. Epilepsy and Behavior Reports. Elsevier Inc.; 2024. doi:10.1016/j.ebr.2024.100651

8. Auvin S, Wirrell E, Donald KA, Berl M, Hartmann H, Valente KD, et al. Systematic review of the screening, diagnosis, and management of ADHD in children with epilepsy. Consensus paper of the Task Force on Comorbidities of the ILAE Pediatric Commission. Epilepsia. 2018 Oct 1;59(10):1867–80. doi:10.1111/epi.14549 PubMed PMID: 30178479.

9. Kilicaslan EE, Türe HS, Kasal Mİ, Çavuş NN, Akyüz DA, Akhan G, et al. Differences in obsessive–compulsive symptom dimensions between patients with epilepsy with obsessive–compulsive symptoms and patients with OCD. Epilepsy and Behavior. 2020 Jan 1;102. doi:10.1016/j.yebeh.2019.106640 PubMed PMID: 31805512.

10. Talıbov T, İnci M, Ismayılov R, Elmas S, Büyüktopçu E, Kepenek AO, et al. The relationship of psychiatric comorbidities and symptoms, quality of life, and stigmatization in patients with epilepsy. Epilepsy and Behavior. 2024 Jul 1;156. doi:10.1016/j.yebeh.2024.109838 PubMed PMID: 38768552.

11. Luoni C, Bisulli F, Canevini MP, De Sarro G, Fattore C, Galimberti CA, et al. Determinants of health-related quality of life in pharmacoresistant epilepsy: Results from a large multicenter study of consecutively enrolled patients using validated quantitative assessments. Epilepsia. 2011 Dec;52(12):2181–91. doi:10.1111/j.1528-1167.2011.03325.x PubMed PMID: 22136077.

12. Taylor RS, Sander JW, Taylor RJ, Baker GA. Predictors of health-related quality of life and costs in adults with epilepsy: A systematic review. Epilepsia. 2011. p. 2168–80. doi:10.1111/j.1528-1167.2011.03213.x PubMed PMID: 21883177.

13. Keezer MR, Sisodiya SM, Sander JW. Comorbidities of epilepsy: Current concepts and future perspectives. The Lancet Neurology. Lancet Publishing Group; 2016. p. 106–15. doi:10.1016/S1474-4422(15)00225-2 PubMed PMID: 26549780.

14. Seidenberg M, Pulsipher DT, Hermann B. Association of epilepsy and comorbid conditions. Future Neurology. 2009. p. 663–8. doi:10.2217/fnl.09.32 PubMed PMID: 20161538.

15. Golyala A, Kwan P. Drug development for refractory epilepsy: The past 25 years and beyond. Seizure. 2017;44:147–56. doi:10.1016/j.seizure.2016.11.022 PubMed PMID: 28017578.

16. Mula M, Kanner AM, Jetté N, Sander JW. Psychiatric Comorbidities in People With Epilepsy. Neurology: Clinical Practice. Lippincott Williams and Wilkins; 2021. p. E112–20. doi:10.1212/CPJ.0000000000000874

17. Galli F, Riquin E, Angers C, Cisneros-Franco FJM, Friedrich F. Psychiatric symptoms and comorbidities in patients with drug-resistant epilepsy in presurgical assessment—A prospective explorative single center study. Report.

18. Shi YQ, Yang HC, He C, Wang YH, Zheng J, Wang XY, et al. Inflammatory links between epilepsy and depression: a review of mechanisms and therapeutic strategies. Frontiers in Neuroscience. Frontiers Media SA; 2025. doi:10.3389/fnins.2025.1614297

19. Basu T, Antonoudiou P, Weiss GL, Coleman EM, David J, Friedman D, et al. Hypothalamic–Pituitary–Adrenal Axis Dysfunction Elevates SUDEP Risk in a Sex-Specific Manner. eNeuro. 2024 Jul 1;11(7). doi:10.1523/ENEURO.0162-24.2024 PubMed PMID: 38914464.

20. Cnops V, Iyer VR, Parathy N, Wong P, Dawe GS. Test, rinse, repeat: A review of carryover effects in rodent behavioral assays. Neuroscience and Biobehavioral Reviews. Elsevier Ltd; 2022. doi:10.1016/j.neubiorev.2022.104560 PubMed PMID: 35124156.

21. Wiltschko AB, Johnson MJ, Iurilli G, Peterson RE, Katon JM, Pashkovski SL, et al. Mapping Sub-Second Structure in Mouse Behavior. Neuron. 2015;88(6):1121–35. doi:10.1016/j.neuron.2015.11.031 PubMed PMID: 26687221.

22. Markowitz J, Gillis W, Beron C, Neufeld S, Robertson K, Bhagat N, et al. The striatum organizes 3D behavior via moment-to-moment action selection. Cell. 2018;174(1):44–58. doi:10.1007/978-3-319-09244-7_5

23. Wiltschko AB, Tsukahara T, Zeine A, Anyoha R, Gillis WF, Markowitz JE, et al. Revealing the structure of pharmacobehavioral space through motion sequencing. Nat Neurosci. 2020 Nov 1;23(11):1433–43. doi:10.1038/s41593-020-00706-3 PubMed PMID: 32958923.

24. Markowitz JE, Gillis WF, Jay M, Wood J, Harris RW, Cieszkowski R, et al. Spontaneous behaviour is structured by reinforcement without explicit reward. Nature. 2023;614(7946):108–17. doi:10.1038/s41586-022-05611-2 PubMed PMID: 36653449.

25. Gschwind T, Zeine A, Raikov I, Markowitz JE, Gillis WF, Felong S, et al. Hidden behavioral fingerprints in epilepsy. Neuron. 2023;111(9):1440–1452.e5. doi:10.1016/j.neuron.2023.02.003 PubMed PMID: 36841241.

26. Silva RR, Munoz DM, Alpert M. Carbamazepine Use in Children and Adolescents with Features of Attention-Deficit Hyperactivity Disorder: A Meta-Analysis. J Am Acad Child Adolesc Psychiatry. 1996;35(3):352–8. doi:10.1097/00004583-199603000-00017 PubMed PMID: 8714324.

27. Grunze A, Amann BL, Grunze H. Efficacy of carbamazepine and its derivatives in the treatment of bipolar disorder. Medicina (Lithuania). 2021 May 1;57(5). doi:10.3390/medicina57050433 PubMed PMID: 33946323.

28. Weisler RH. Carbamazepine extended-release capsules in bipolar disorder. Neuropsychiatric Disease and Treatment. 2006. Report.

29. Hagberg AA, Schult DA, Swart PJ. Exploring Network Structure, Dynamics, and Function using NetworkX [Internet]. 2008. Report. Available from: http://conference.scipy.org/proceedings/SciPy2008/paper_2

30. Hoffman OR, Koehler JL, Ezekiel J, Espina C, Patterson AM, Gohar ES, et al. Disease modification upon 2 weeks of tofacitinib treatment in a mouse model of chronic epilepsy. Sci. Transl. Med [Internet]. 2025. Report. Available from: https://www.science.org

31. Birbeck GL, Hays RD, Cui X, Vickrey BG. Seizure reduction and quality of life improvements in people with epilepsy. Epilepsia. 2002 May 24;43(5):535–8. doi:10.1046/j.1528-1157.2002.32201.x PubMed PMID: 12027916.

32. Siebenbrodt K, Willems LM, von Podewils F, Mross PM, Strüber M, Langenbruch L, et al. Determinants of quality of life in adults with epilepsy: a multicenter, cross-sectional study from Germany. Neurol Res Pract. 2023;5(1). doi:10.1186/s42466-023-00265-5

33. Sadleir LG, Scheffer IE. Automatisms in Absence Seizures in Children With Idiopathic Generalized Epilepsy. Report.

34. Yang B, Mo J, Zhang C, Wang X, Sang L, Zheng Z, et al. Clinical features of automatisms and correlation with the seizure onset zones: A cluster analysis of 74 surgically-treated cases. Seizure. 2022 Jan 1;94:82–9. doi:10.1016/j.seizure.2021.11.015 PubMed PMID: 34872021.

35. Bird JS, Shah E, Shotbolt P. Epilepsy and concomitant obsessive–compulsive disorder. Epilepsy Behav Case Rep. 2018 Jan 1;10:106–10. doi:10.1016/j.ebcr.2018.07.001

36. Rogers DC, Dittner AJ, Rimes KA, Chalder T. Fatigue in an adult attention deficit hyperactivity disorder population: A trans-diagnostic approach. British Journal of Clinical Psychology. 2017 Mar 1;56(1):33–52. doi:10.1111/bjc.12119 PubMed PMID: 27918087.

37. Barkmeier DT, Senador D, Leclercq K, Pai D, Hua J, Boutros NN, et al. Electrical, molecular and behavioral effects of interictal spiking in the rat. Neurobiol Dis. 2012 Jul;47(1):92–101. doi:10.1016/j.nbd.2012.03.026 PubMed PMID: 22472188.

38. Kanemura H. Association between epilepsy and attention deficit/hyperactivity disorder – correlation between interictal epileptiform discharges and behavioral disturbances. Brain and Development. Elsevier B.V.; 2025. doi:10.1016/j.braindev.2025.104403

39. Silvestri R, Gagliano A, Calarese T, Aricò I, Cedro C, Condurso R, et al. Ictal and interictal EEG abnormalities in ADHD children recorded over night by video-polysomnography. Epilepsy Res. 2007 Jul;75(2–3):130–7. doi:10.1016/j.eplepsyres.2007.05.007 PubMed PMID: 17588723.

40. Bakke KA, Larsson PG, Eriksson AS, Eeg-Olofsson O. Levetiracetam reduces the frequency of interictal epileptiform discharges during NREM sleep in children with ADHD. European Journal of Paediatric Neurology. 2011 Nov;15(6):532–8. doi:10.1016/j.ejpn.2011.04.014 PubMed PMID: 21683631.

41. Goos G, Hartmanis J, Van J, Board LE, Hutchison D, Kanade T, et al. Network Analysis: Methodological Foundations. Lecture Notes in. Heidelberg; 1973. Report. doi:10.1007/b106453

42. Ding Q, Zhang F, Feng Y, Wang H. Carbamazepine restores neuronal signaling, protein synthesis, and cognitive function in a mouse model of fragile X syndrome. Int J Mol Sci. 2020 Dec 1;21(23). doi:10.3390/ijms21239327 PubMed PMID: 33297570.

43. Joffe RT, Swinson RP. Carbamazepine in Obsessive-Compulsive Disorder. 1987. Report.

44. Egashira N, Abe M, Shirakawa A, Niki T, Mishima K, Iwasaki K, et al. Effects of mood stabilizers on marble-burying behavior in mice: Involvement of GABAergic system. Psychopharmacology (Berl). 2013 Mar;226(2):295–305. doi:10.1007/s00213-012-2904-9 PubMed PMID: 23086022.

45. Gierbolini J, Giarratano M, Benbadis SR. Carbamazepine-related antiepileptic drugs for the treatment of epilepsy-a comparative review. Expert Opinion on Pharmacotherapy. Taylor and Francis Ltd; 2016. p. 885–8. doi:10.1517/14656566.2016.1168399 PubMed PMID: 26999402.

