## supplemental figures for "A network approach with motion sequencing reveals hidden patterns of repetitive behavior in a pre-clinical model of epilepsy" (from another version of this preprint: version 2, 2026-04-08)

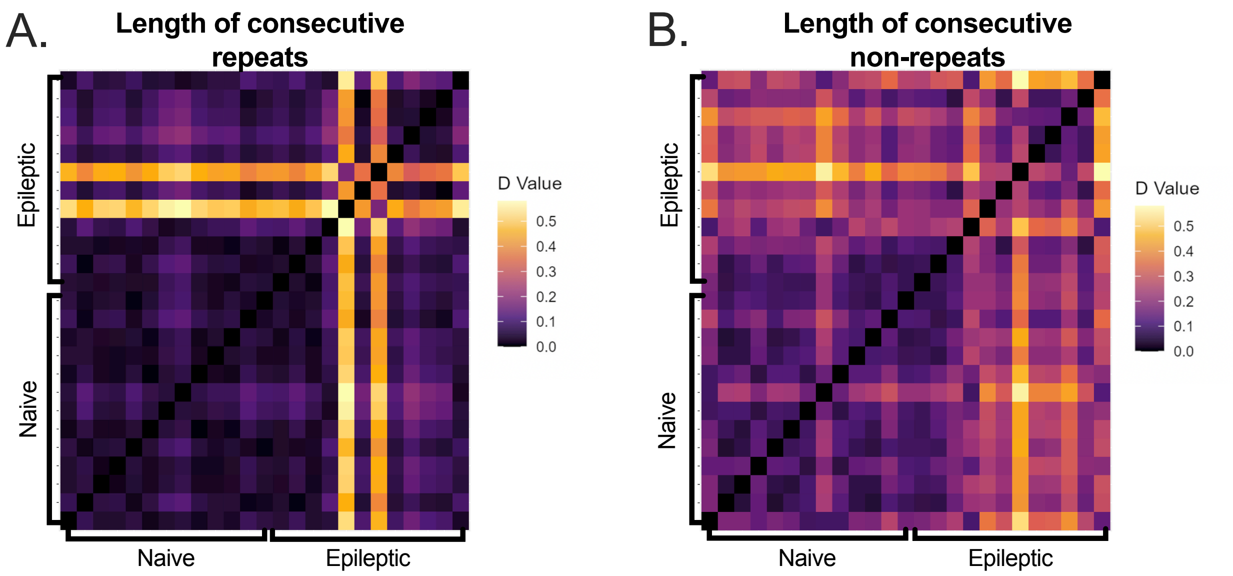


**Fig. S1. PERMANOVA heatmaps for length of consecutive repeats and length of consecutive non-repeats.**
Heatmaps depict the Kolmogorov-Smirnov distance for each possible pairing for all mice across both conditions, epileptic and naïve, for both **(A)** length of consecutive repeats and **(B)** length of consecutive non-repeats.

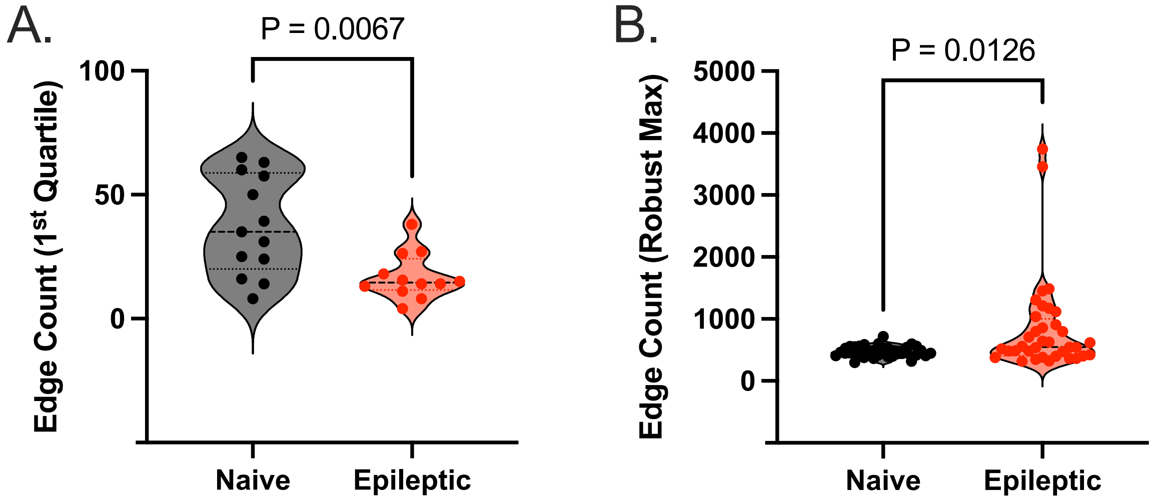


**Fig. S2. Epilepsy leads to a reduction in edge count in the first quartile and an increase at the robust max.**
Violin plots depict the edge counts for naïve and epileptic mice at the **(A)** 1st quartile (p = 0.0067) and **(B)** the robust max (p = 0.0126). The robust max is calculated using the top three values. Each point represents a mouse. Statistical analysis conducted using the Mann-Whitney test.

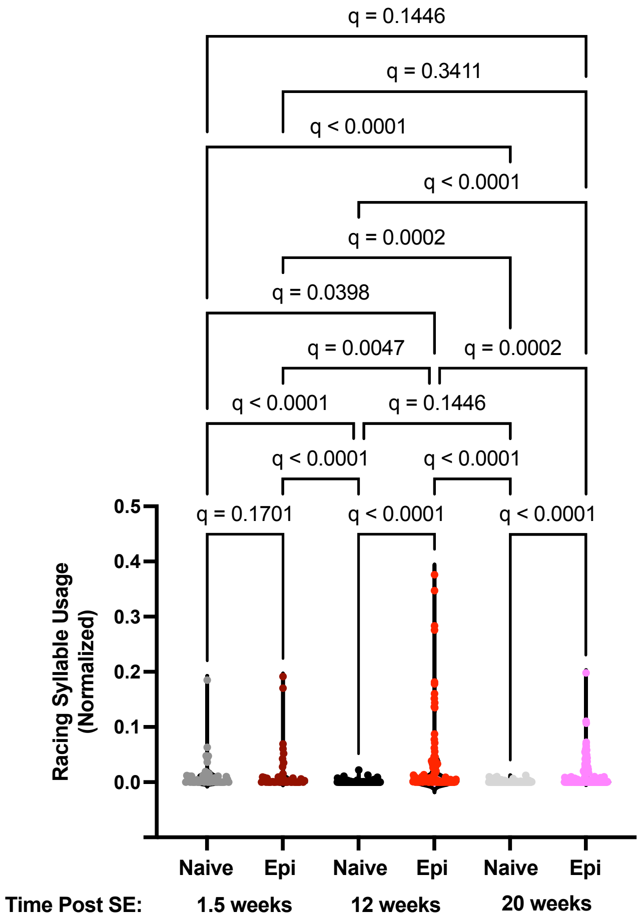


**Fig. S3. Racing syllable usage increases as epilepsy progresses.**
Violin plots depict the racing syllable usage for naïve and epileptic mice at three different timepoints (1.5-, 12-, and 20-weeks post SE). Each syllable was normalized to the total number of syllables in each respective syllable list. Each point represents a racing syllable. Statistical analysis conducted using the Kruskal-Wallis test with BKY post-hoc.

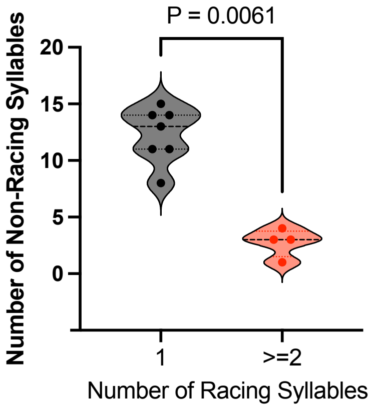


**Fig. S4. Mice with multiple racing syllables display use less non-racing syllables.**
Violin plots depict the number of non-racing syllables with a normalized usage over 0.025 for epileptic mice 12 weeks post SE ($p = 0.0061$). Each syllable was normalized to the total number of syllables in each respective syllable list. Each point represents a mouse. Statistical analysis conducted using the Mann-Whitney Test.

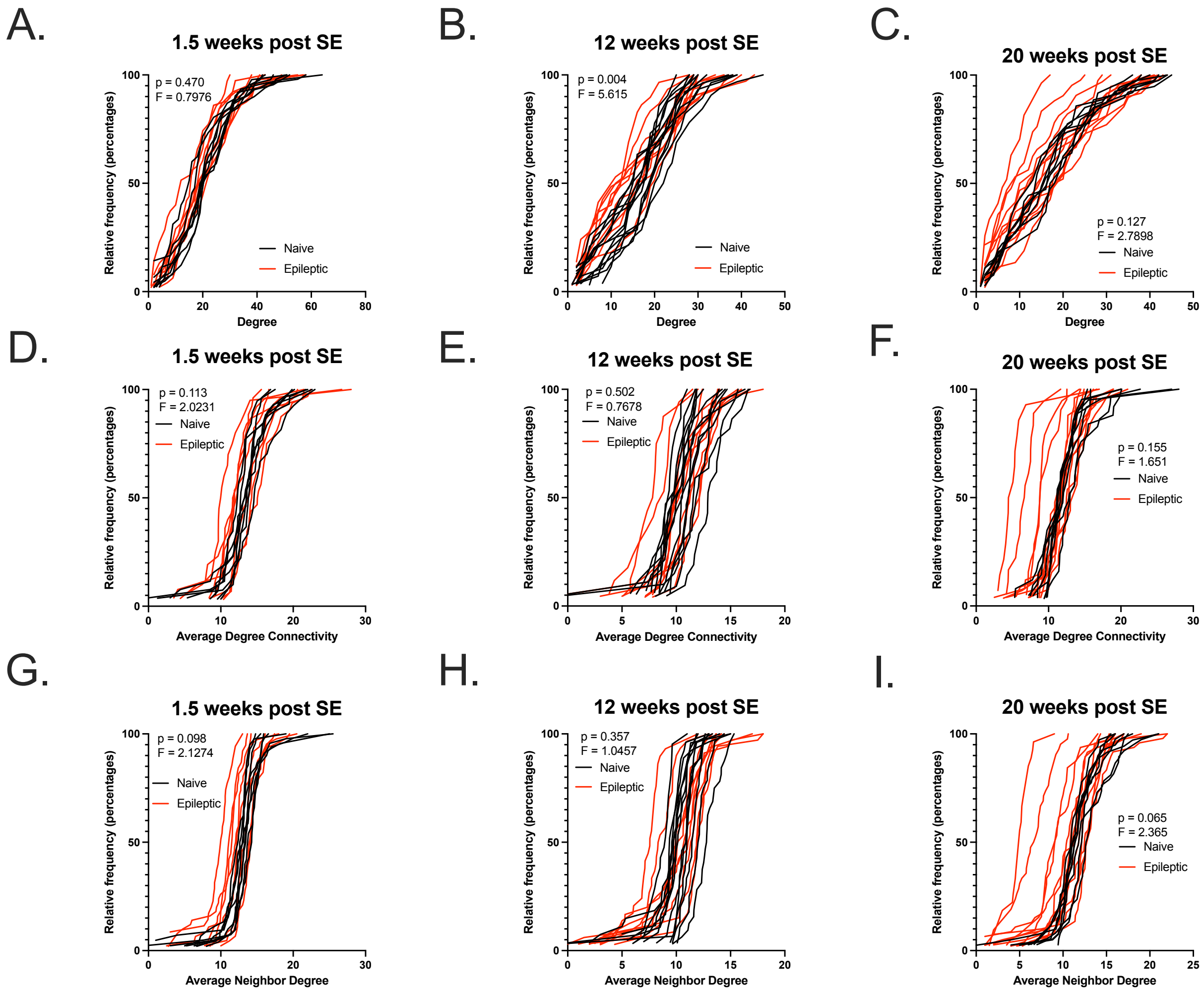


**Fig. S5. Analysis of degree, average degree connectivity, and average neighbor degree throughout disease progression.**

Network metrics for 3 different time points (1.5-, 12-, and 20-weeks post SE) displayed a cumulative frequency distribution for each condition, naïve and epileptic. Degree at **(A)** 1.5-weeks post SE (Pseudo-F = 0.7976, p = 0.470), **(B)** 12-weeks post SE (Pseudo-F = 5.615, p = 0.004), and **(C)** 20-weeks post SE (Pseudo-F = 2.7898, p = 0.127). Average degree connectivity at **(D)** 1.5-weeks post SE (Pseudo-F = 2.0231, p = 0.113), **(E)** 12-weeks post SE (Pseudo-F =0.7678, p = 0.502), and **(F)** 20-weeks post SE (Pseudo-F = 1.561, p = 0.155). Average neighbor degree at **(G)** 1.5-weeks post SE (Pseudo-F = 2.1274, p = 0.096), **(H)** 12-weeks post SE (Pseudo-F = 1.0457, p = 0.357), and **(I)** 20-weeks post SE (Pseudo-F = 2.365, p = 0.065). Each line represents a mouse. All statistics performed using a PERMANOVA (999 permutations) based on the Kolmogorov-Smirnov distance.

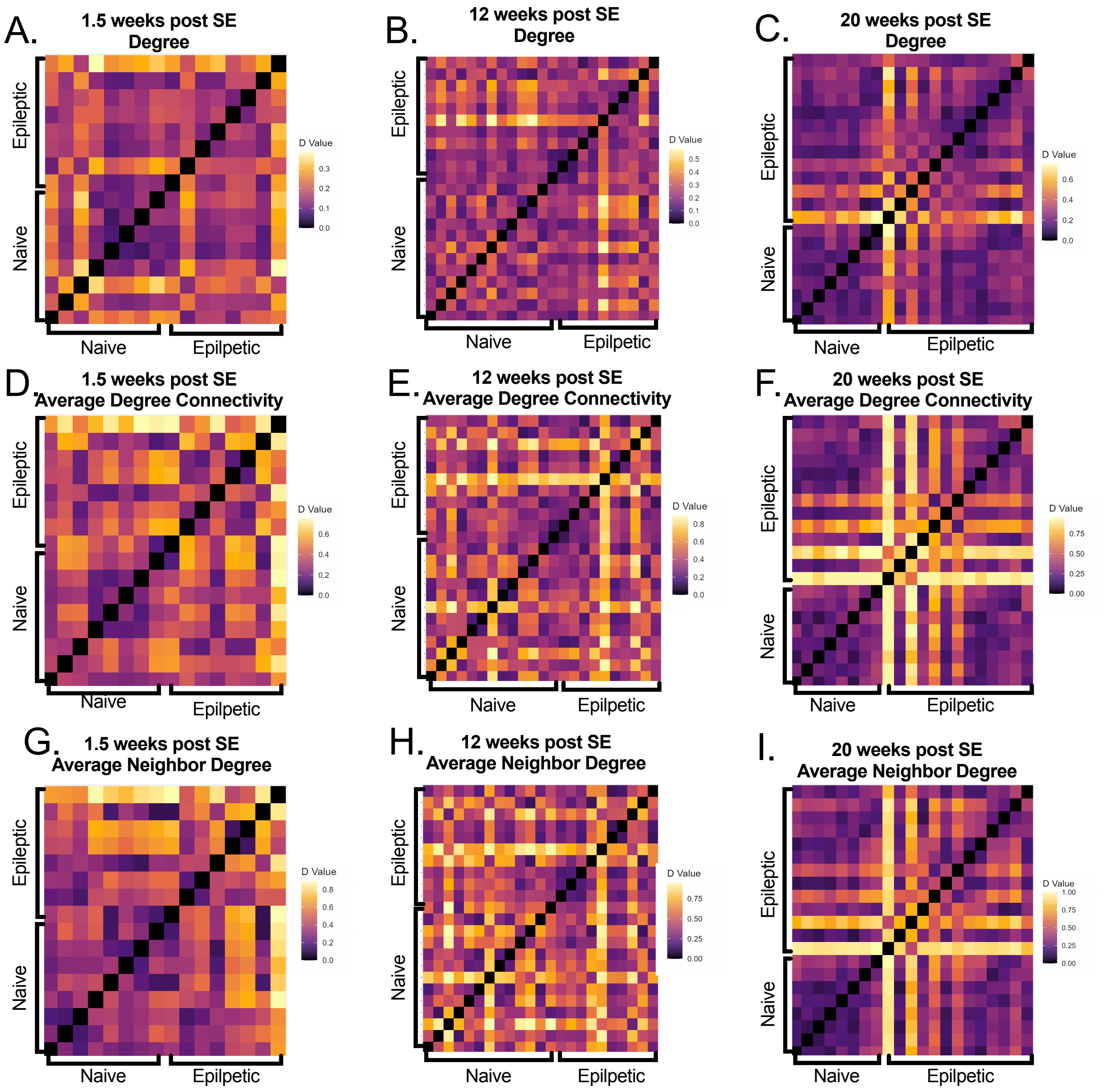


**Fig. S6. PERMANOVA heatmaps for degree, average degree connectivity, and average neighbor degree during disease progression.**

Heatmaps depict the Kolmogorov-Smirnov distance for each possible pairing for all mice across both conditions, epileptic and naïve, at 3 different timepoints (1.5-, 12-, and 20-weeks post SE) in for **(A)** degree, **(B)** average degree connectivity, and **(C)** average neighbor degree.

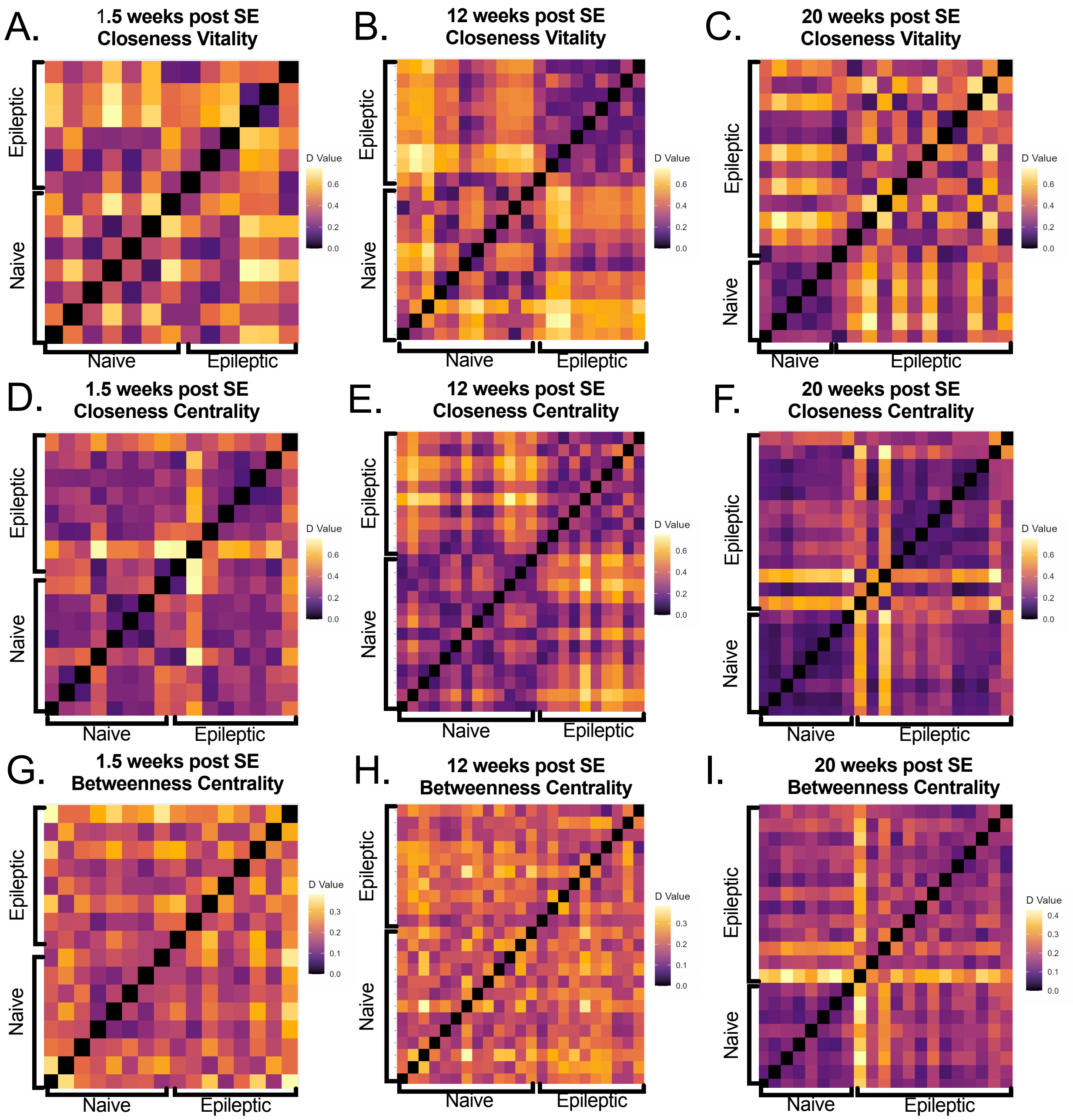


**Fig. S7. PERMANOVA heatmaps for closeness vitality, closeness centrality, and betweenness centrality during disease progression.**
Heatmaps depict the Kolmogorov-Smirnov distance for each possible pairing for all mice across both conditions, epileptic and naïve, at 3 different timepoints (1.5-, 12-, and 20-weeks post SE) in for **(A)** closeness vitality, **(B)** closeness centrality, and **(C)** betweenness centrality.

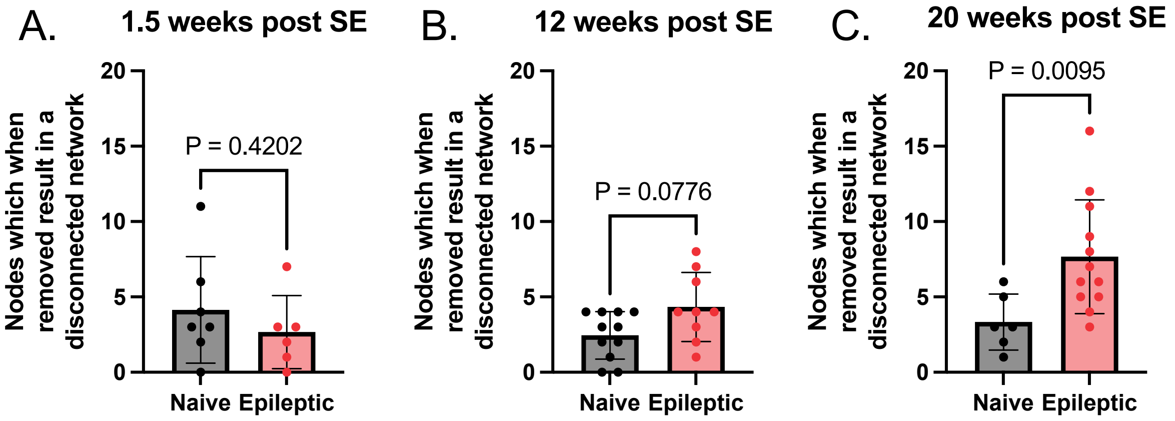


**Fig. S8. The number of nodes essential for network connectivity is higher in epileptic vs naïve mice at 20-weeks post SE.**
Bar graphs depict the number of nodes which when removed results in a disconnected network for **(A)** 1.5-, **(B)** 12-, and **(C)** 20-weeks post SE. All statistics performed using a Mann-Whitney test.

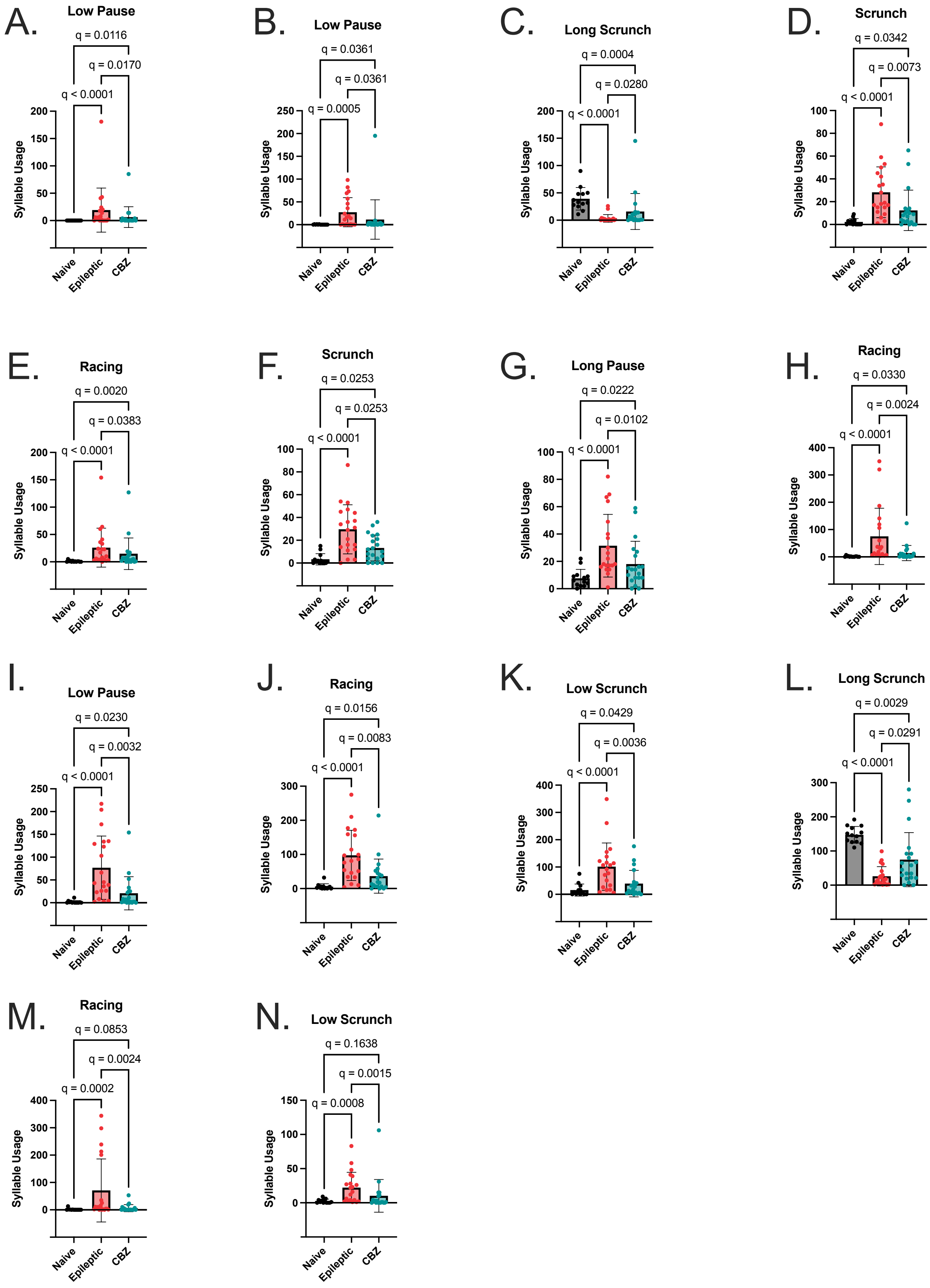
A.
Low Pause
q = 0.0116
q = 0.0170
q < 0.0001
Syllable Usage
Naive
Epileptic
CBZ
B.
Low Pause
q = 0.0361
q = 0.0361
q = 0.0005
C.
Long Scrunch
q = 0.0004
q = 0.0280
q < 0.0001
D.
Scrunch
q = 0.0342
q = 0.0073
q < 0.0001
E.
Racing
q = 0.0020
q = 0.0383
q < 0.0001
F.
Scrunch
q = 0.0253
q = 0.0253
q < 0.0001
G.
Long Pause
q = 0.0222
q = 0.0102
q < 0.0001
H.
Racing
q = 0.0330
q = 0.0024
q < 0.0001
I.
Low Pause
q = 0.0230
q = 0.0032
q < 0.0001
J.
Racing
q = 0.0156
q = 0.0083
q < 0.0001
K.
Low Scrunch
q = 0.0429
q = 0.0036
q < 0.0001
L.
Long Scrunch
q = 0.0029
q = 0.0291
q < 0.0001
M.
Racing
q = 0.0853
q = 0.0024
q = 0.0002
N.
Low Scrunch
q = 0.1638
q = 0.0015
q = 0.0008

**Fig. S9. CBZ tends to rescue syllables upregulated during epilepsy.** Bar graphs depict syllable usage for naïve, epileptic, and CBZ treated mice. **(A)** Low pause (naïve and epileptic, $q <0.0001$) (naïve and CBZ, $q = 0.0116$) (epileptic and CBZ, $q = 0.0170$). **(B)** Low Pause (naïve and epileptic, $q = 0.0005$) (naïve and CBZ, $q = 0.0361$) (epileptic and CBZ, $q = 0.0361$). **(C)** Long scrunch (naïve and epileptic, $q < 0.0001$) (naïve and CBZ, $q = 0.0004$) (epileptic and CBZ, $q = 0.0280$). **(D)** Scrunch (naïve and epileptic, $q < 0.0001$) (naïve and CBZ, $q = 0.0342$) (epileptic and CBZ, $q = 0.0073$). **(E)** Racing (naïve and epileptic, $q < 0.0001$) (naïve and CBZ, $q = 0.0020$) (epileptic and CBZ, $q = 0.0383$). **(F)** Scrunch (naïve and epileptic, $q < 0.0001$) (naïve and CBZ, $q = 0.0253$) (epileptic and CBZ, $q = 0.0253$). **(G)** Long Pause (naïve and epileptic, $q < 0.0001$) (naïve and CBZ, $q = 0.0222$) (epileptic and CBZ, $q = 0.0102$). **(H)** Racing (naïve and epileptic, $q < 0.0001$) (naïve and CBZ, $q = 0.0330$) (epileptic and CBZ, $q = 0.0024$). **(I)** Low Pause (naïve and epileptic, $q < 0.0001$) (naïve and CBZ, $q = 0.0230$) (epileptic and CBZ, $q = 0.0032$). **(J)** Racing (naïve and epileptic, $q < 0.0001$) (naïve and CBZ, $q = 0.0156$) (epileptic and CBZ, $q = 0.00083$). **(K)** Low Scrunch (naïve and epileptic, $q < 0.0001$) (naïve and CBZ, $q = 0.0429$) (epileptic and CBZ, $q = 0.0036$). **(L)** Long Scrunch (naïve and epileptic, $q < 0.0001$) (naïve and CBZ, $q = 0.0029$) (epileptic and CBZ, q 0.0291). **(M)** Racing (naïve and epileptic, $q = 0.0002$) (naïve and CBZ, $q = 0.0853$) (epileptic and CBZ, $q = 0.0024$). **(N)** Low Scrunch (naïve and epileptic, $q = 0.0008$) (naïve and CBZ, $q = 0.1638$) (epileptic and CBZ, $q = 0.0015$). Each point represents a mouse. All statistics performed using Kruskal-Wallis test with BKY post hoc.

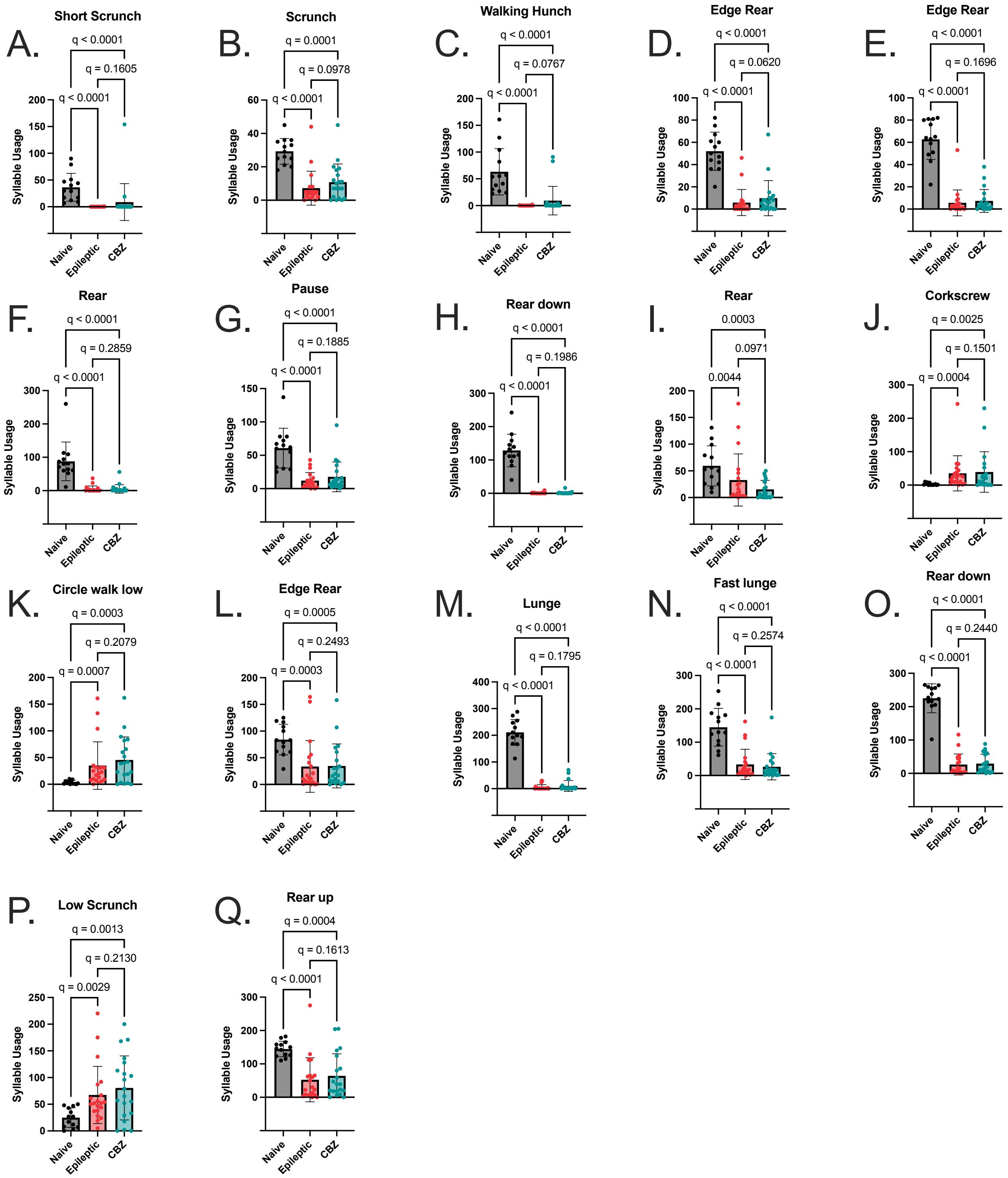
A.
Short Scrunch
q < 0.0001
q = 0.1605
q < 0.0001
Syllable Usage
Naive
Epileptic
CBZ
B.
Scrunch
q < 0.0001
q = 0.0978
q < 0.0001
C.
Walking Hunch
q < 0.0001
q = 0.0767
q < 0.0001
D.
Edge Rear
q < 0.0001
q = 0.0620
q < 0.0001
E.
Edge Rear
q < 0.0001
q = 0.1696
q < 0.0001
F.
Rear
q < 0.0001
q = 0.2859
q < 0.0001
G.
Pause
q < 0.0001
q = 0.1885
q < 0.0001
H.
Rear down
q < 0.0001
q = 0.1986
q < 0.0001
I.
Rear
0.0003
0.0971
0.0044
J.
Corkscrew
q = 0.0025
q = 0.1501
q = 0.0004
K.
Circle walk low
q = 0.0003
q = 0.2079
q = 0.0007
L.
Edge Rear
q = 0.0005
q = 0.2493
q = 0.0003
M.
Lunge
q < 0.0001
q = 0.1795
q < 0.0001
N.
Fast lunge
q < 0.0001
q = 0.2574
q < 0.0001
O.
Rear down
q < 0.0001
q = 0.2440
q < 0.0001
P.
Low Scrunch
q = 0.0013
q = 0.2130
q = 0.0029
Q.
Rear up
q = 0.0004
q = 0.1613
q < 0.0001

**Fig. S10. CBZ fails to rescue syllables downregulated by epilepsy.** Bar graphs depict syllable usage for naïve, epileptic, and CBZ treated mice. **(A)** Short Scrunch (naïve and epileptic, $q < 0.0001$) (naïve and CBZ, $q < 0.0001$) (epileptic and CBZ, $q = 0.1605$). **(B)** Scrunch (naïve and epileptic, $q < 0.0001$) (naïve and CBZ, $q = 0.0001$) (epileptic and CBZ, $q = 0.0978$). **(C)** Walking Hunch (naïve and epileptic, $q < 0.0001$) (naïve and CBZ, $q < 0.0001$) (epileptic and CBZ, $q = 0.0767$). **(D)** Edge Rear (naïve and epileptic, $q < 0.0001$) (naïve and CBZ, $q < 0.0001$) (epileptic and CBZ, $q = 0.0626$). **(E)** Edge Rear (naïve and epileptic, $q < 0.0001$) (naïve and CBZ, $q < 0.0001$) (epileptic and CBZ, $q = 0.1696$). **(F)** Rear (naïve and epileptic, $q < 0.0001$) (naïve and CBZ, $q < 0.0001$) (epileptic and CBZ, $q = 0.2859$). **(G)** Pause (naïve and epileptic, $q < 0.0001$) (naïve and CBZ, $q < 0.0001$) (epileptic and CBZ, $q = 0.1885$). **(H)** Rear down (naïve and epileptic, $q < 0.0001$) (naïve and CBZ, $q < 0.0001$) (epileptic and CBZ, $q = 0.1986$). **(I)** Rear (naïve and epileptic, $q = 0.0044$) (naïve and CBZ, $q = 0.0003$) (epileptic and CBZ, $q = 0.0971$). **(J)** Corkscrew (naïve and epileptic, $q = 0.0004$) (naïve and CBZ, $q = 0.0025$) (epileptic and CBZ, $q = 0.1501$). **(K)** Circle walk low (naïve and epileptic, $q = 0.0007$) (naïve and CBZ, $q = 0.0003$) (epileptic and CBZ, $q = 0.2079$). **(L)** Edge Rear (naïve and epileptic, $q = 0.0003$) (naïve and CBZ, $q = 0.0005$) (epileptic and CBZ, $q = 0.2493$). **(M)** Lunge (naïve and epileptic, $q < 0.0001$) (naïve and CBZ, $q < 0.0001$) (epileptic and CBZ, $q = 0.1795$). **(N)** Fast Lunge (naïve and epileptic, $q < 0.0001$) (naïve and CBZ, $q < 0.0001$) (epileptic and CBZ, $q = 0.2574$). **(O)** Rear down (naïve and epileptic, $q < 0.0001$) (naïve and CBZ, $q < 0.0001$) (epileptic and CBZ, $q = 0.2440$). **(P)** Low Scrunch (naïve and epileptic, $q = 0.0029$) (naïve and CBZ, $q = 0.0013$) (epileptic and CBZ, 0.2130). **(Q)** Rear up (naïve and epileptic, $q < 0.0001$) (naïve and CBZ, $q = 0.0004$) (epileptic and CBZ, $q = 0.1613$). Each point represents a mouse. All statistics performed using Kruskal-Wallis test with BKY post hoc.

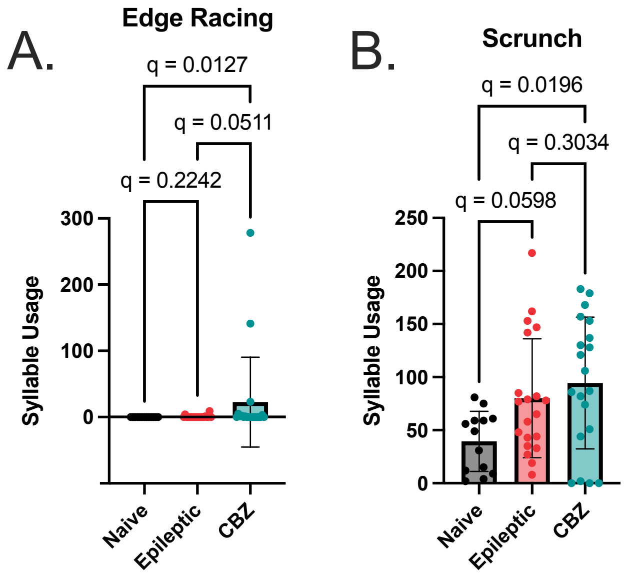


**Fig. S11. CBZ treatment leads to off target behavioral effects.** Bar graphs depict syllable usage for naïve, epileptic, and CBZ treated mice. **(A)** Edge Racing (naïve and epileptic, q = 0.2242) (naïve and CBZ, q = 0.0127) (epileptic and CBZ, q = 0.0511). **(B)** Scrunch (naïve and epileptic, q = 0.0598) (naïve and CBZ, q = 0.096) (epileptic and CBZ, q = 0.3034). Each point represents a mouse. All statistics performed using Kruskal-Wallis test with BKY post hoc.

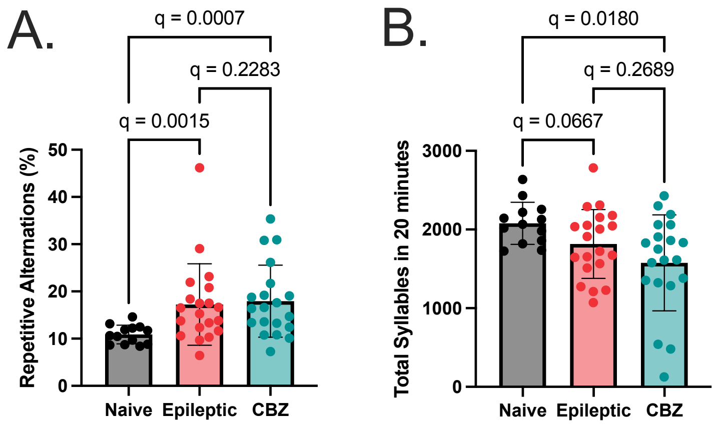


**Fig. S12. CBZ treatment reduces the total number of syllables traveled in 20 minutes.** Bar graphs depict **(A)** the percent repetitive alternations (naïve and epileptic, q = 0.0015) (naïve and CBZ, q = 0.0007) (epileptic and CBZ, q = 0.2283) and **(B)** total syllables in 20 minutes (naïve and epileptic, q = 0.0667) (naïve and CBZ, q = 0.0180) (epileptic and CBZ, q = 0.2689). Each point representing a mouse. All statistics performed using Kruskal-Wallis test with BKY post hoc.

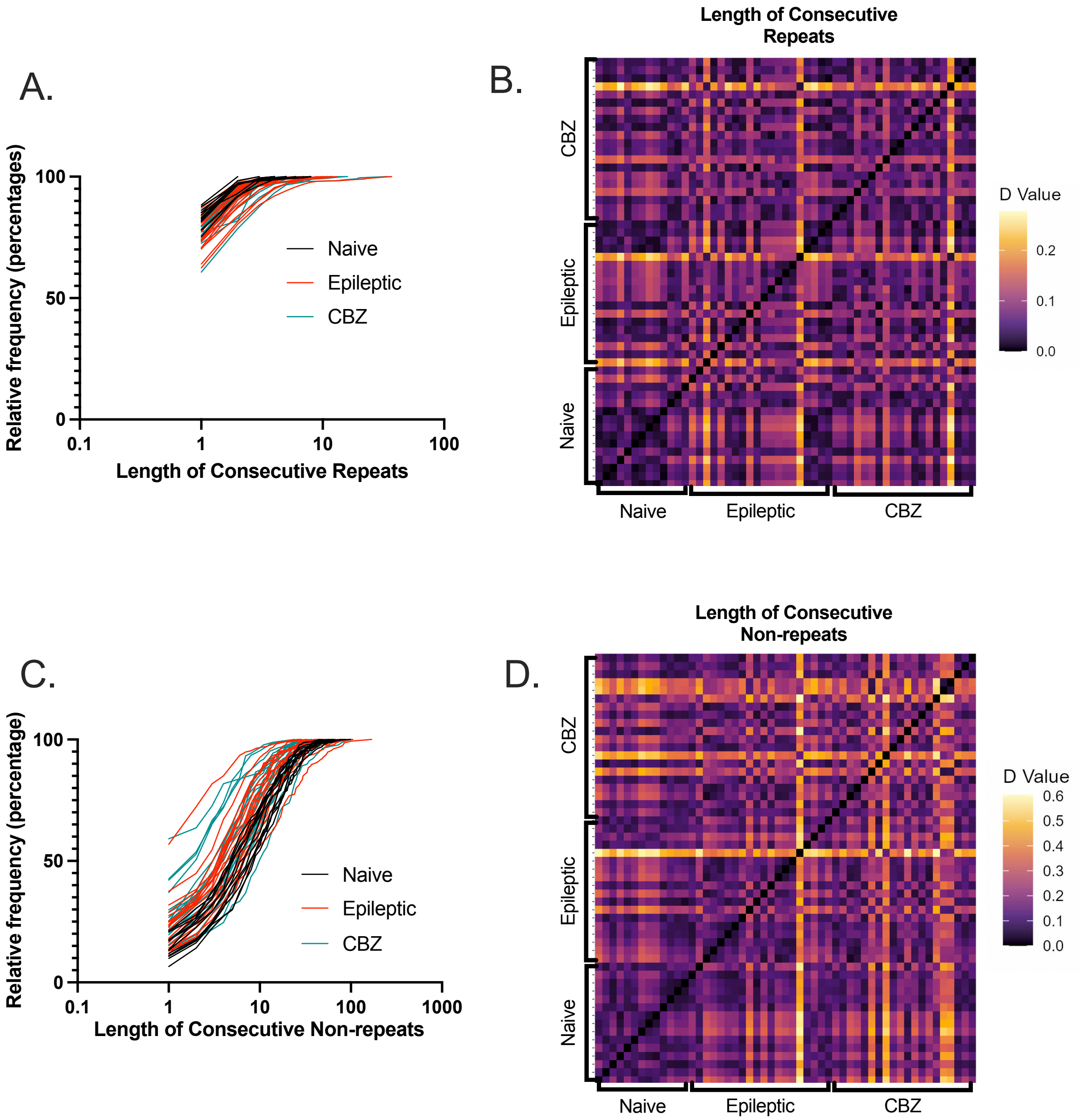


**Fig. S13. CBZ does not rescue length of consecutive repeats and non-repeats in epileptic mice. (A)**Cumulative of the length of consecutive repetitive alternations for naïve, epileptic and CBZ treated mice. (naïve and epileptic, Pseudo-F = 7.5431, p = 0.012) (naïve and CBZ, Pseudo-F = 9.091, p = 0.003) (epileptic and CBZ, Pseudo-F = -0.1854, p = 0.933). Each line represents a mouse. **(B)** A heatmap depicts the Kolmogorov-Smirnov distance for each possible pairing for all mice across naïve, epileptic and CBZ treated mice. **(C)** Cumulative of the length of consecutive non-repetitive alterations for epileptic and naïve mice (naïve and epileptic, Pseudo-F = 7.5363, p = 0.006) (naïve and CBZ, Pseudo-F = 8.3961, p = 0.006) (epileptic and CBZ, Pseudo-F = 0.2873, p = 0.774). Each line represents a mouse. **(D)** A heatmap depicts the Kolmogorov-Smirnov distance for each possible pairing for all mice across naïve, epileptic and CBZ treated mice. All statistics performed using a PERMANOVA (999 permutations) based on the Kolmogorov-Smirnov distance and a Benjamini-Hochberg post hoc.

A.

B.

C.

D.

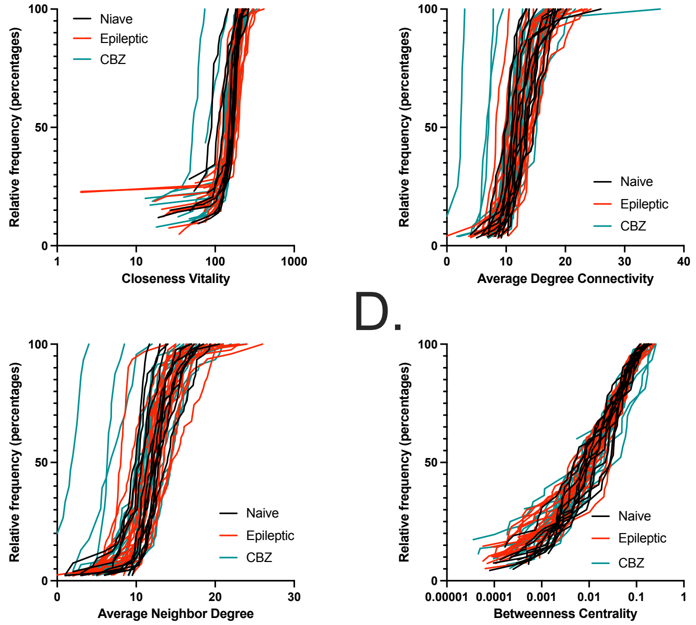


**Fig. S14. Analysis of closeness vitality, average degree connectivity, average neighbor degree, and betweenness centrality in CBZ treated mice.**
Network metrics for 3 different conditions (naïve, epileptic, and CBZ treated) with each line representing a mouse. **(A)** Closeness vitality (naïve and epileptic, Pseudo-F = 0.6542, p = 0.789) (naïve and CBZ, Pseudo-F = 0.4501, p = 0.789) (epileptic and CBZ, Pseudo-F = 0.8752, p = 0.789). **(B)** Average degree connectivity (naïve and epileptic, Pseudo-F = 0.2918, p = 0.844) (naïve and CBZ, Pseudo-F = 0.6576, p = 0.844) (epileptic and CBZ, Pseudo-F = 0.9184, p = 0.844). **(C)** Average neighbor degree (naïve and epileptic, Pseudo-F = 0.9763, p = 0.566) (naïve and CBZ, Pseudo-F = 0.733, q = 0.566) (epileptic and CBZ, Pseudo-F = 0.9414, p = 0.566). **(D)** Betweenness centrality (naïve and epileptic, Pseudo-F = 1.5026, p = 0.5085) (naïve and CBZ, Pseudo-F = 1.381, p = 0.5085) (epileptic and CBZ, Pseudo-F = 0.7274, p = 0.632). All statistics performed using a PERMANOVA (999 permutations) based on the Kolmogorov-Smirnov distance and a Benjamini-Hochberg post hoc.

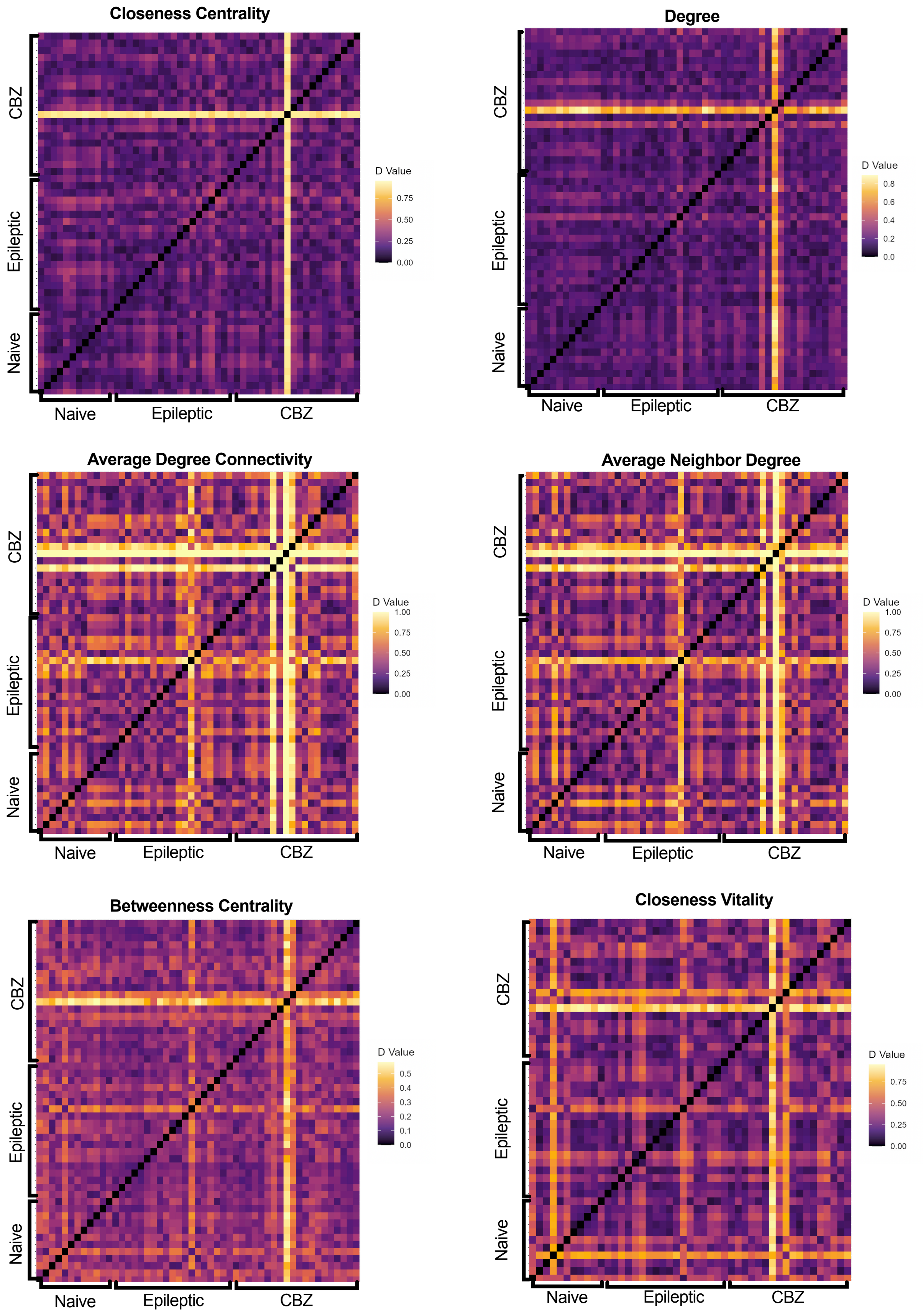
Closeness Centrality
CBZ
Epileptic
Naive
Naive
Epileptic
CBZ
D Value
0.75
0.50
0.25
0.00
Degree
CBZ
Epileptic
Naive
Naive
Epileptic
CBZ
D Value
0.8
0.6
0.4
0.2
0.0
Average Degree Connectivity
CBZ
Epileptic
Naive
Naive
Epileptic
CBZ
D Value
1.00
0.75
0.50
0.25
0.00
Average Neighbor Degree
CBZ
Epileptic
Naive
Naive
Epileptic
CBZ
D Value
1.00
0.75
0.50
0.25
0.00
Betweenness Centrality
CBZ
Epileptic
Naive
Naive
Epileptic
CBZ
D Value
0.5
0.4
0.3
0.2
0.1
0.0
Closeness Vitality
CBZ
Epileptic
Naive
Naive
Epileptic
CBZ
D Value
0.75
0.50
0.25
0.00

**Fig. S15. PERMANOVA heatmaps for network metrics for CBZ experiment.** Heatmaps the Kolmogorov-Smirnov distance for each possible pairing for all mice across both conditions, epileptic and naïve, at 3 different conditions (naïve, epileptic, and CBZ treated) in for **(A)** closeness centrality, **(B)** degree, **(C)** average degree connectivity, **(D)** average neighbor degree, **(E)** betweenness centrality, and **(F)** closeness vitality.

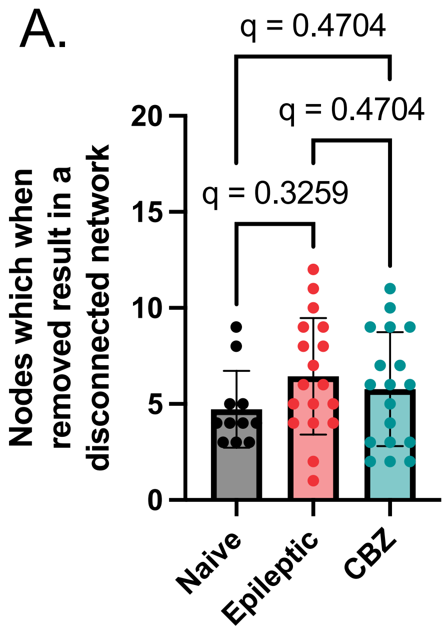


**Fig. S16. The number of nodes essential for network connectivity is not different across naïve, epileptic, and CBZ treated mice.**
Bar graphs depict the number of nodes which when removed results in a disconnected network for **(A)** naïve, epileptic, and CBZ treated mice. All statistics performed using a Mann-Whitney test.

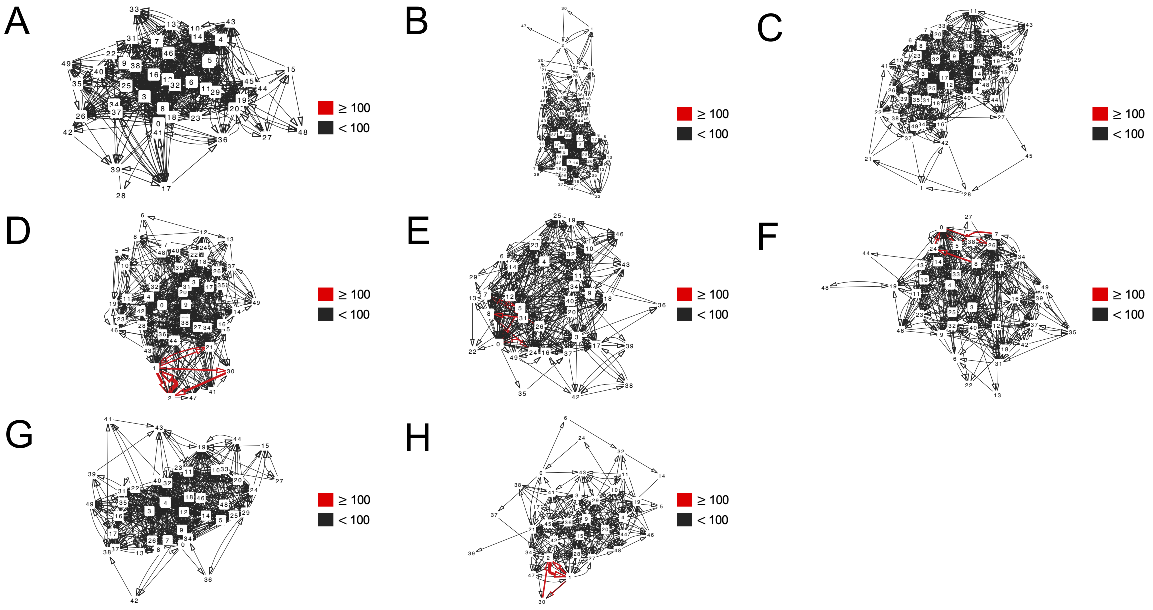


**Fig. S17. 1.5 weeks post SE epileptic mice individual behavioral networks**
Directed networks wherein each node represents a syllable, and each edge represents a transition. The edge is determined by the number of transitions. Edges with a frequency greater than or equal to 100 are colored red. Networks are ordered by mouse from **(A)** Epileptic 1 to **(H)** Epileptic 8.

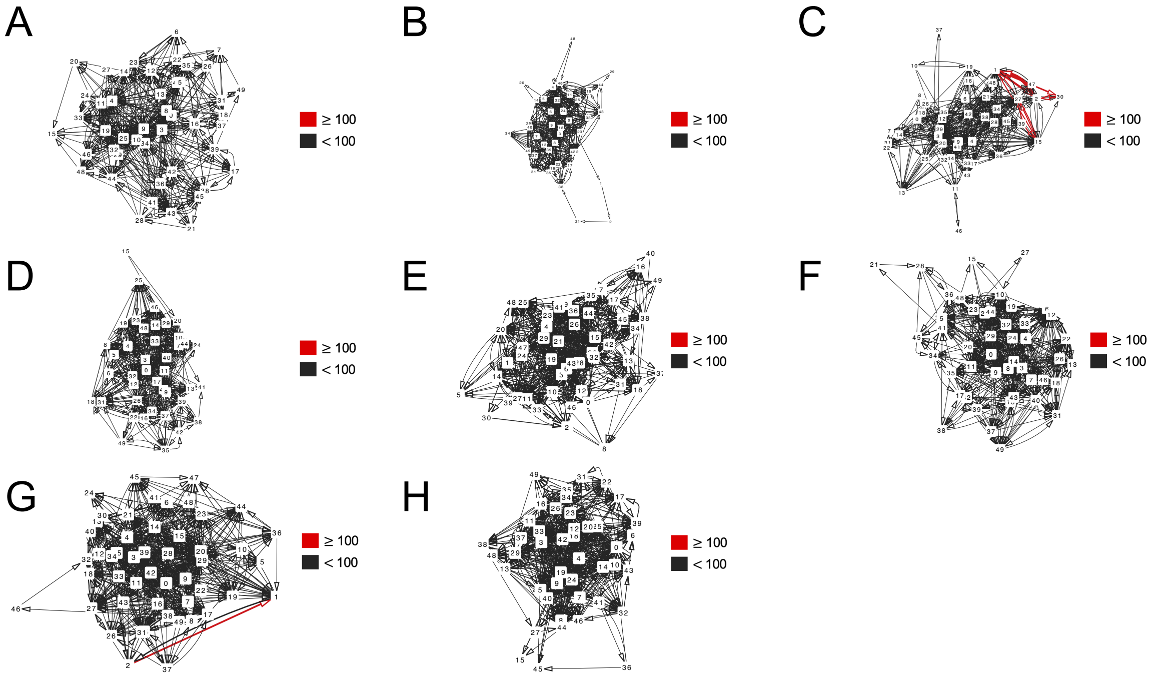


**Fig. S18. 1.5 weeks post SE age-matched naive mice individual behavioral networks**
Directed networks wherein each node represents a syllable, and each edge represents a transition. The edge is determined by the number of transitions. Edges with a frequency greater than or equal to 100 are colored red. Networks are ordered by mouse from **(A)** Naive 1 to **(H)** Naive 8.

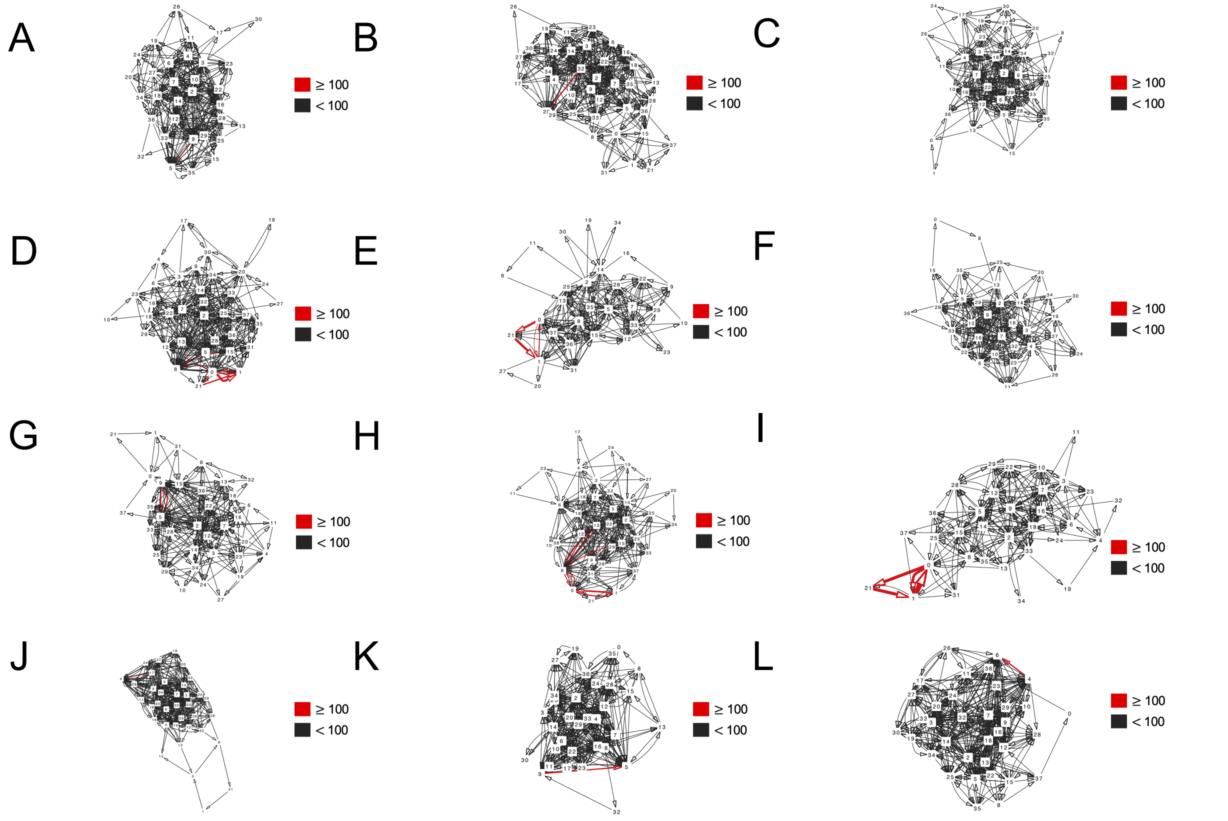


**Fig. S19. 15 weeks post SE epileptic mice individual behavioral networks**
Directed networks wherein each node represents a syllable, and each edge represents a transition. The edge is determined by the number of transitions. Edges with a frequency greater than or equal to 100 are colored red. Networks are ordered by mouse from **(A)** Epileptic 1 to **(L)** Epileptic 12.

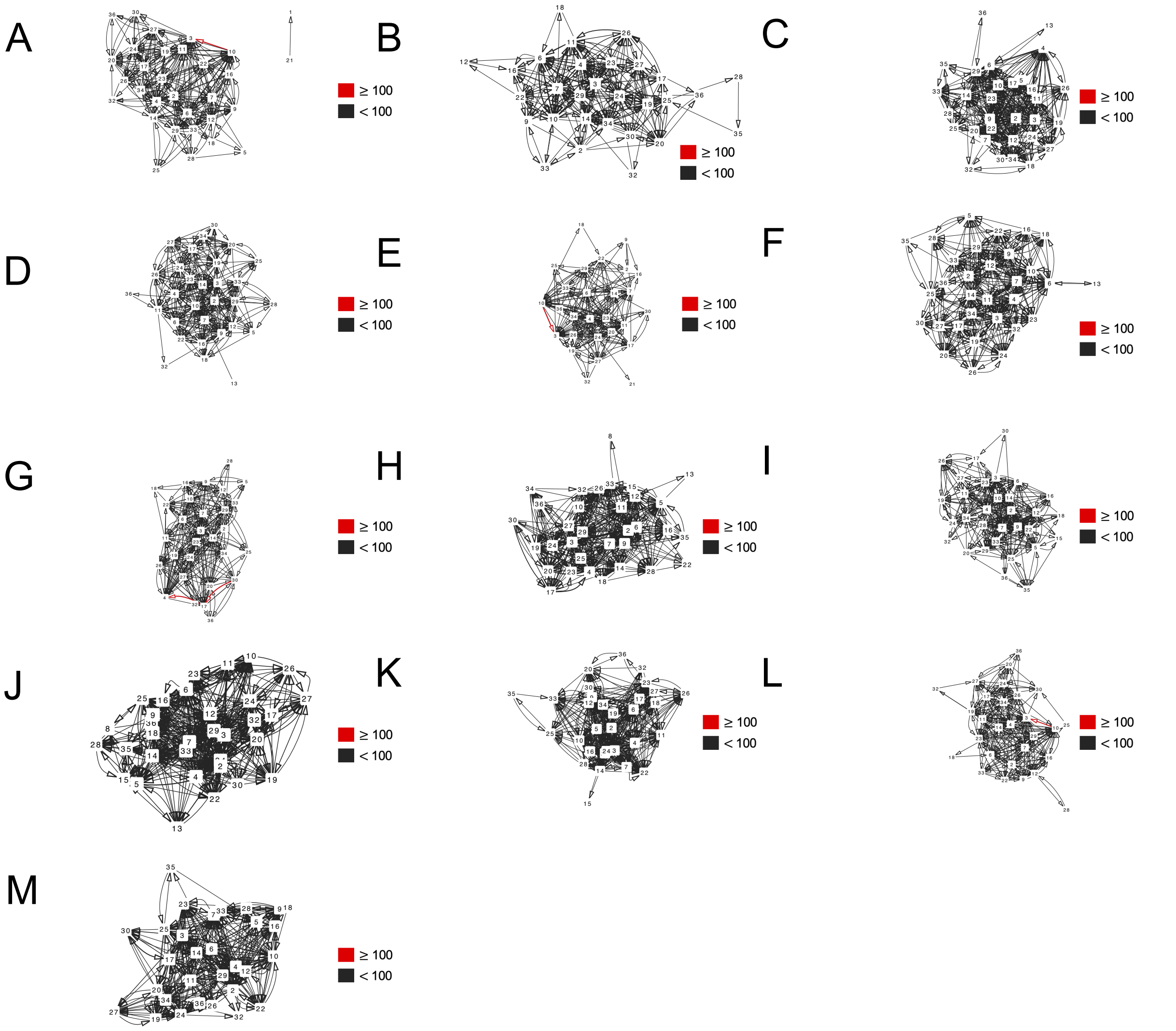


**Fig. S20. 12 weeks post SE aged-matched naive mice individual behavioral networks**
Directed networks wherein each node represents a syllable, and each edge represents a transition. The edge is determined by the number of transitions. Edges with a frequency greater than or equal to 100 are colored red. Networks are ordered by mouse from **(A)** Naive 1 to **(M)** Naive 13.

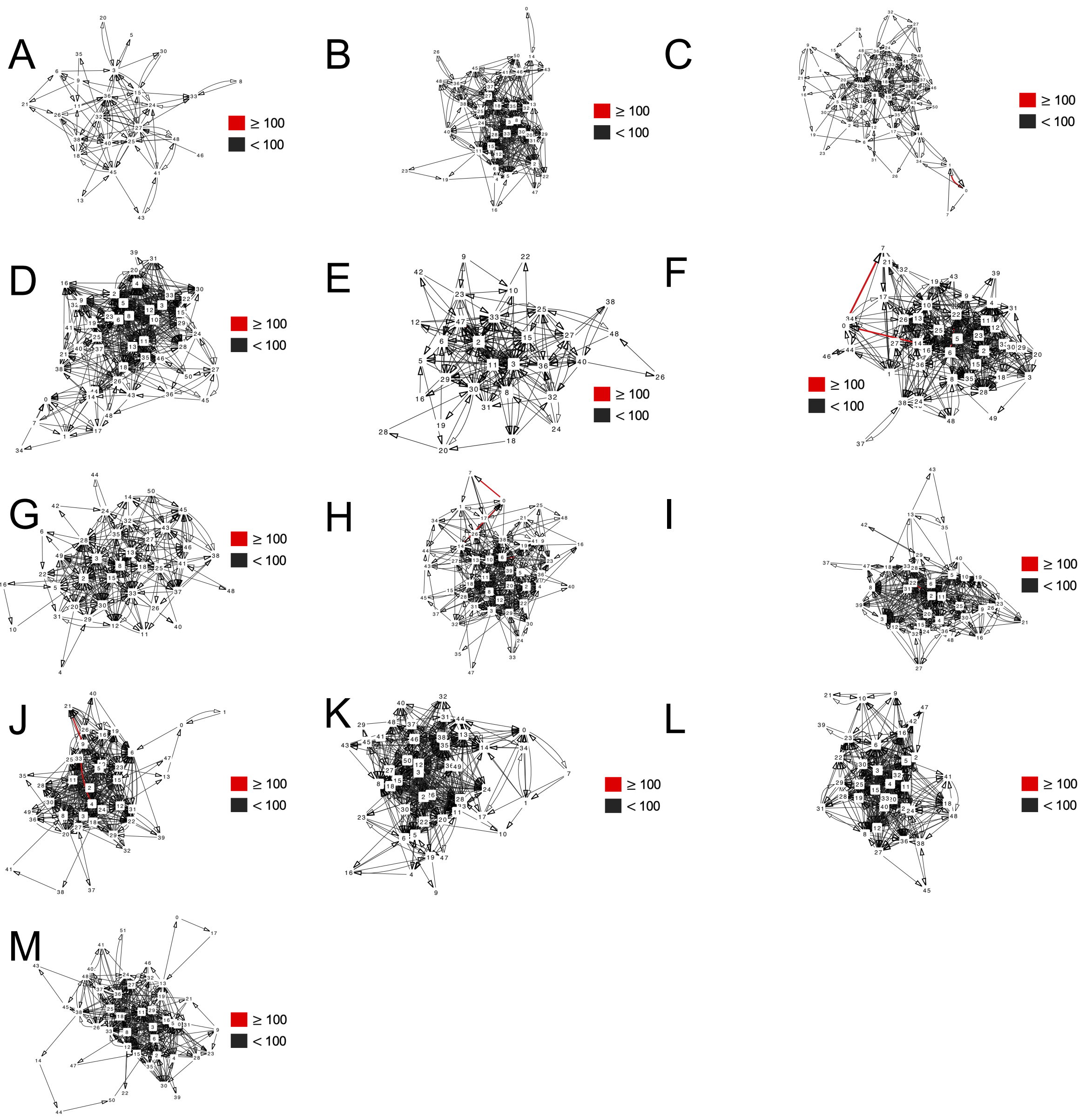


**Fig. S21. 20 weeks post SE epileptic individual behavioral networks**
Directed networks wherein each node represents a syllable, and each edge represents a transition. The edge is determined by the number of transitions. Edges with a frequency greater than or equal to 100 are colored red. Networks are ordered by mouse from **(A)** Epileptic 1 to **(M)** Epileptic 13.

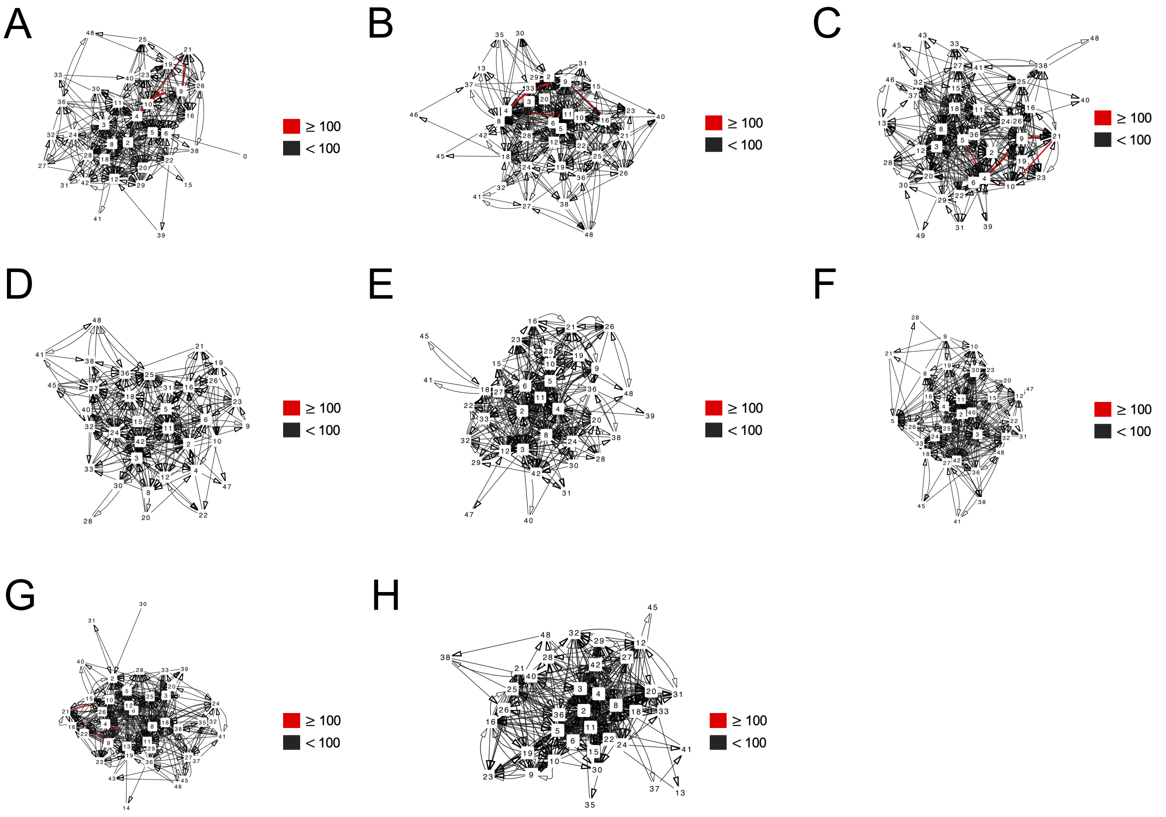


**Fig. S22. 20 weeks post SE age-matched naive individual behavioral networks**
Directed networks wherein each node represents a syllable, and each edge represents a transition. The edge is determined by the number of transitions. Edges with a frequency greater than or equal to 100 are colored red. Networks are ordered by mouse from **(A)** Naive 1 to **(H)** Naïve 8.

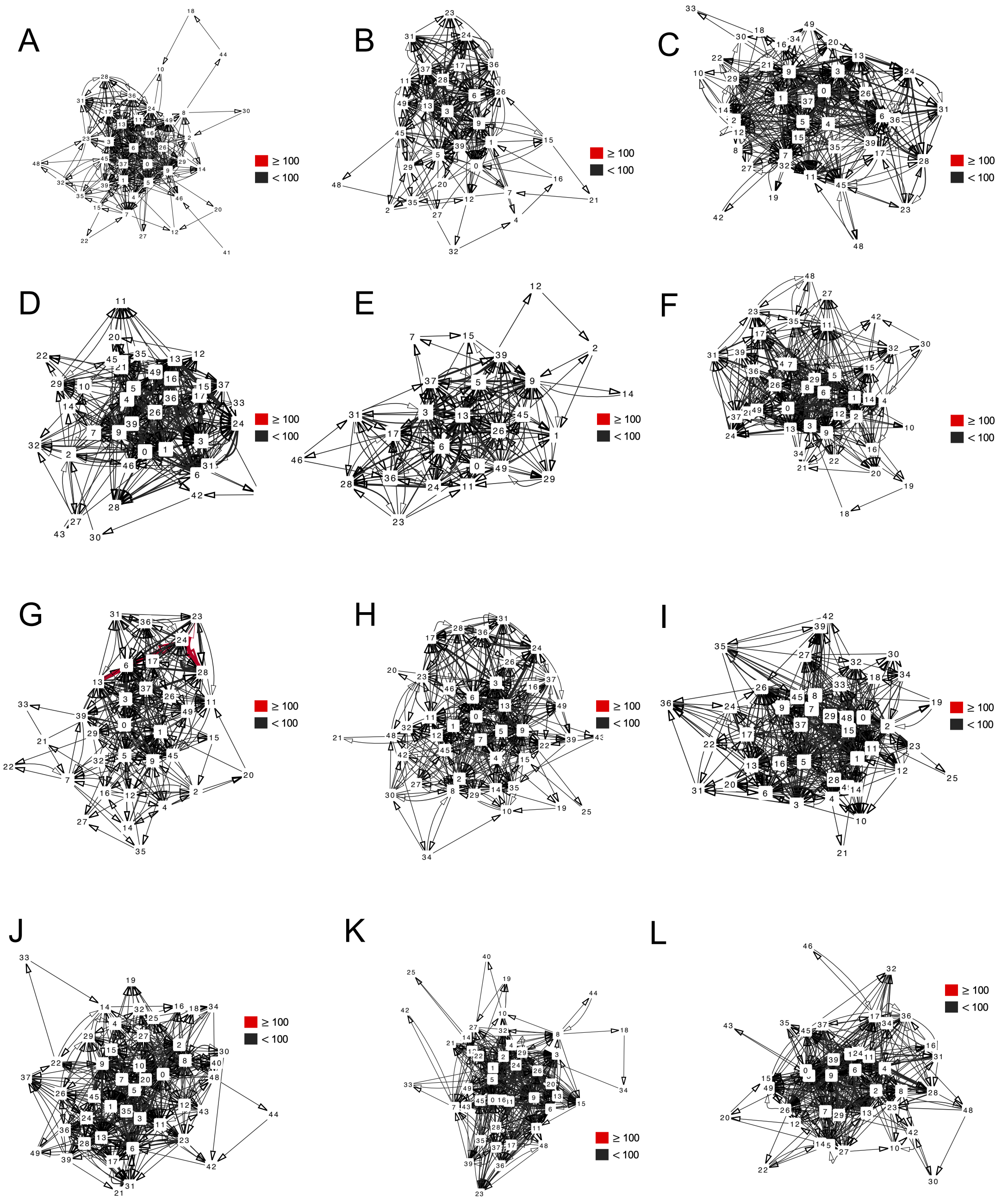


**Fig. S23. 12 weeks post SE age-matched naive individual behavioral networks for CBZ experiment** Directed networks wherein each node represents a syllable, and each edge represents a transition. The edge is determined by the number of transitions. Edges with a frequency greater than or equal to 100 are colored red. Networks are ordered by mouse from **(A)** Naive 1 to **(L)** Naïve 12.

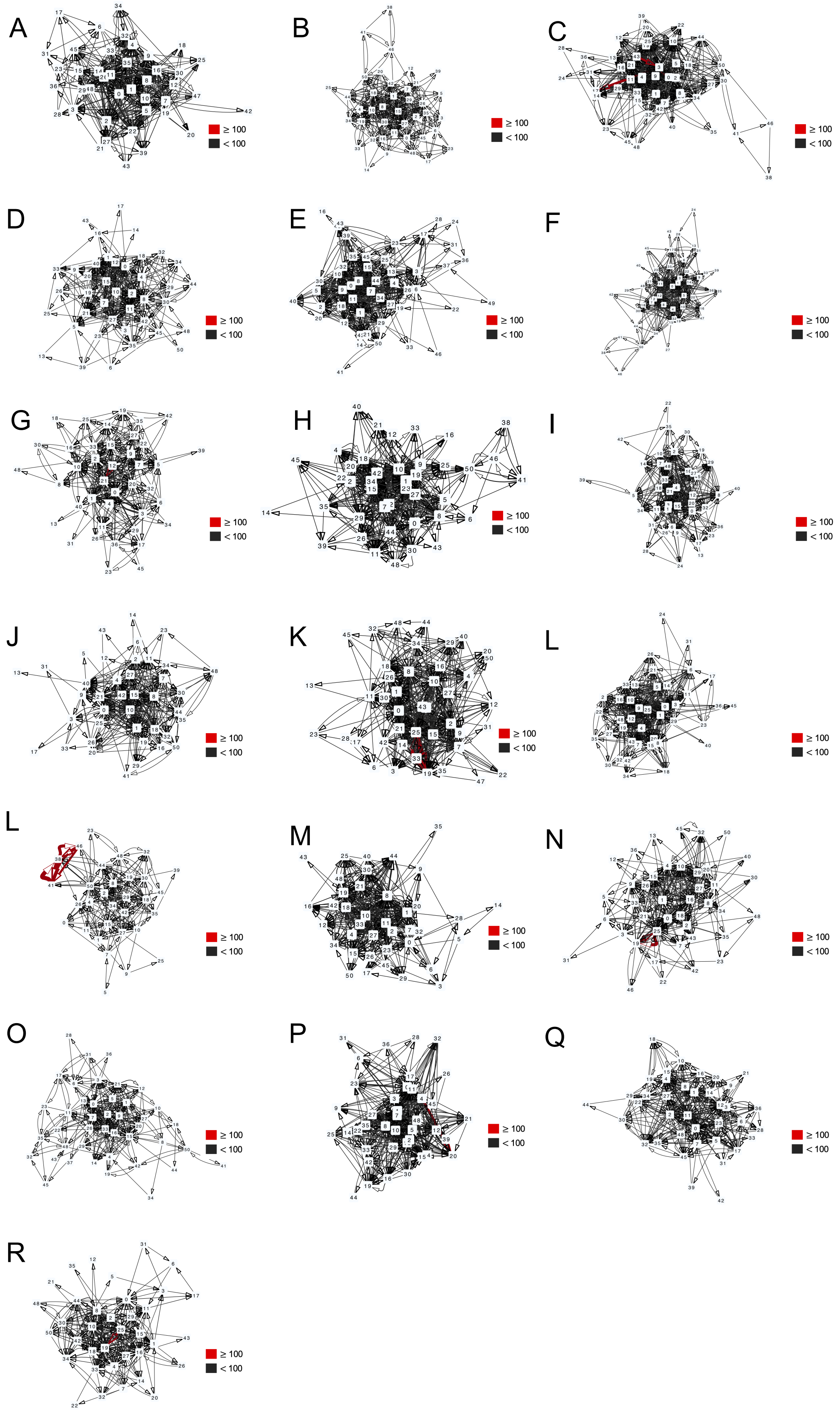
A
B
C
D
E
F
G
H
I
J
K
L
L
M
N
O
P
Q
R
≥ 100
< 100

**Fig. S24. 12 weeks post SE epileptic individual behavioral networks for CBZ experiment**
Directed networks wherein each node represents a syllable, and each edge represents a transition. The edge is determined by the number of transitions. Edges with a frequency greater than or equal to 100 are colored red. Networks are ordered by mouse from **(A)** Epileptic 1 to **(R)** Naïve 18.

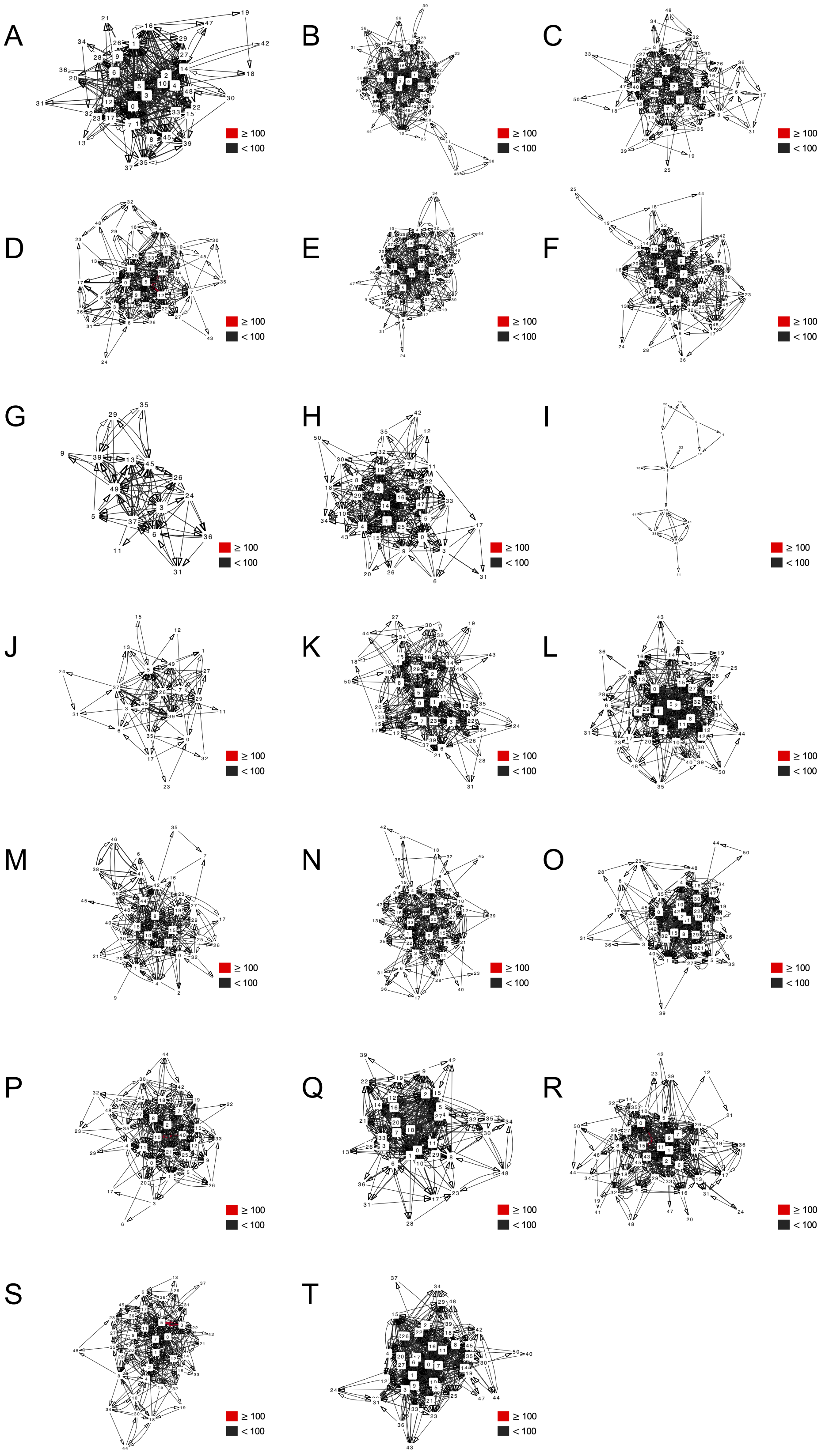
A
B
C
D
E
F
G
H
I
J
K
L
M
N
O
P
Q
R
S
T
≥ 100
< 100

**Fig. S25. 12 weeks post SE CBZ treated individual behavioral networks for CBZ experiment**
Directed networks wherein each node represents a syllable, and each edge represents a transition. The edge is determined by the number of transitions. Edges with a frequency greater than or equal to 100 are colored red. Networks are ordered by mouse from **(A)** Epileptic 1 to **(T)** Naïve 20.

| | Participated | Failed to Participate | Fisher's Exact Test p-value |
|---|---|---|---|
| Age matched Naive | 9 | 0 | |
| Epileptic (34 weeks post SE) | 2 | 6 | 0.0023 |

**Table S1. Participation in the spontaneous alternation Y maze test is impaired 34 weeks post SE in epileptic mice.** Participation is classified as making 15 or more alternations in the Y maze in 8 minutes. Statistical analysis performed using the Fisher's exact test.
